# Wheels within wheels, Nested Function-value Traits as a Tool for Modeling Ontogeny

**DOI:** 10.1101/2021.03.16.435729

**Authors:** John G. Hodge, Andrew N. Doust

## Abstract

Plant morphologies exhibit a wide array of outcomes that have evolved as a consequence adapting to a wide array of ecological pressures. These disparate morphologies have provided a rich field for comparative morphologists, developmental biologists, and geneticists to explore. Ultimately the array of variation observed in nature across different plant species is built on the same functional unit, the phytomer, which is composed of a leaf, a node, and an internode. Sequentially produced phytomers exhibit heteroblasty, that is, a gradual or abrupt change in shape, either due to size changes or changes due to reproductive phase. The progression of shape change over time is often indirectly measured by sampling several stages of plant growth and comparing allometric relationships between shape variables. However, a more precise method is to use an absolute time scale and measure shape change of sequential organs directly. In this study we use such time-dependent measurements to build a general model of organ growth for several Setaria genotypes, for both leaves and internodes. We term this the second-order function-value trait (2FVT) model, because it generalizes individual function-value trait models generated for each organ. This model reduces phenotypic noise by averaging the general trend of ontogeny and provides a quantitative tool to describe where and when phenotypic shifts occur during the ontogenies of different genotypes. The ability to recognize how ontogenetic variation is distributed within equivalent positions of the body plan at the interspecific level can be used as a tool to explore various questions related to growth and form in plants both for comparative morphology and developmental genetics.

## Introduction

Plant vegetative architectures are ultimately the product of iterative developmental decisions governing growth. These iterative choices are made through the periodic production of phytomers, sequential organ units composed of a leaf, internode, and lateral bud. Over a plant’s life history, the modification of phytomers based on specific physio-developmental roles gives rise to the wide range of morphologies observed in plant lineages. The derivation of the constituent organs formed within these iterative subunits along an axis, particularly the leaves, bracts, and floral organs, has been a topic that has intrigued botanists for centuries. Best known among these is Johann Wolfgang von Goethe who focused many of his theories on the “metamorphosis” of lateral organs via ontogenetic plasticity (Arber, 1946). There is a rich body of literature which documents the various processes at work to drive these morphological shifts. Variation can be either differences in growth over time, within and between ontogenetic trajectories of different genetic backgrounds, and/or shifts in growth strategy.

Comparisons of organs can be done between those on the same plant (an ontogenetic series) or between organs of different plants. Anatomical and morphological differences related to ontogeny have also been termed heteroblastic differences, originally used for marked differences in form between juvenile and reproductive stages (Jones 1999, Zotz et al. 2011), but now used as a broad term describing any morphological heterogeneity (Jones 1999). Ontogenetic differences on a single plant have also been termed heteroauxesis (Huxley et al. 1941), and Niklas (1994) makes a sharp distinction between such differences and those that occur on different plants (termed allomorphosis). Like Niklas, we regard it as important to first understand ontogenetic differences before making between genotype comparisons.

The role of time in assessing ontogenetic changes is not always appreciated. Much of the consideration of differences throughout ontogeny has been based on allometric relationships, often gained through destructive sampling of plant growth at only a few timepoints. Allometry establishes the relationship between different shape parameters through time, such as length and breadth of leaf blades, and the time axis can only be constructed in a relative sense (e.g. Sinnott 1921, Hammond et al. 1941, Baker et al. 2015, Chitwood et al. 2015, Chitwood et al. 2017). However, present advances in phenomics via high throughput imagery can provide an absolute time scale for ontogenetic change, and will lead to much increased precision in comparing between ontogenies (e.g. McCormick et al. 2016, Pound et al. 2017, Miao et al. 2020). There is likely untapped potential for combining the strategies of regressing measures against time with the assumption of state change models assumed in either the heteroblasty or allometry literature (Jones 1999, Nijhout & German 2012). An analogous simulation was able to readily resolve subtle variations in branch topology in inflorescences by assuming phase changes between primary and subordinate axes (Prusinkiewicz et al. 2007).

Although they are generally targeted to a specific organ or trait, time-series based approaches have used sigmoidal curves to model organ growth, due to their ability to describe growth that starts slow, undergoes a rapid acceleration, before gradually attenuating as the organ achieves a mature size (Niklas 1994, Nijhout & German, 2012). Moreover, the parameters of these sigmoidal models can be extracted as function-value traits (FVTs), that can be analyzed to reveal the underlying genetic control of these growth processes (Baker et al. 2015). However, studies have tended to concentrate on particular organs and ignore the ontogenetic relationship between organs, although we would expect that models underlying the growth of sequential structures such as leaves should reveal similar fundamental growth processes uncovered by the analysis of FVTs. Using all members of an ontogenetic series to predict the growth characteristics of individual members is a way to reduce noise resulting from measurement error or environmental effects, and thus improve function-value trait comparisons.

In this study we explore whether taking account of the relationship between sequentially produced organs improves prediction and accuracy of models for organ growth. We refer to such an approach as a second-order function value trait model (2FVT model), consisting of a nested regression analysis which attempts to characterize whole plant phenotypes as a phytomeric progression by using a series of sigmoidal models with linked parameters, that are estimated by their position within the plant body. Treating the models in this manner can drastically reduce the overall number of parameters needed to describe observed trait variation for ontogenetic series of organs and improve prediction for values for individual organs.

## Methods

### Collection of time-series data set

The Setaria model system, consisting of the wild species *Setaria viridis* (green foxtail) and the domesticated lineage *Setaria italica* (foxtail millet), as well as several recombinant inbred lines (RIL) from a mapping population created from a cross between domesticated *S. italica* acc. B100 and wild *S. viridis* acc. A10 (Mauro-Herrera et al. 2016). Ten replicates of the two parents, A10 and B100, and three RILs, RIL39, RIL110, and RIL159 (corresponding to RIL32, RIL85, and RIL129, in Feldman et al. 2017) were selected for this study. These were chosen as representative of phenotypic variation in the RIL population for plant height, flowering, leaf stature, and branching. Due to parallax errors in imaging resulting from torsion of the main axis two replicates of RIL110 and one replicate each of RIL39 and RIL159 were omitted from the final pseudo-landmark and morphometric analysis.

Plants were grown under 12-hour day conditions with temperatures varying between 22-28°C during dark and light cycles respectively. Phenotyping was conducted through the use of a Raspberry Pi 3 (model B, version 2) imaging station over the first 30 days of growth, at which point all genotypes except for B100 had flowered (Fig. 1). The first 4 days of growth were omitted from imaging because none of the genotypes had grown out of the pot sufficiently to measure until day 5, and because the first two leaves are embryonic in origin rather than produced by the shoot apical meristem. We used a semi-automated pipeline to create a sparse dataset of pseudo-landmarks that identified leaf tips and ligules (*Acute* workflow) before using PCA and clustering techniques to identify homology groups of the same organ in different images (*Constella* workflow) (Hodge et al. 2021). These homology groups were then annotated with standardized names that reflected their identities as structures of biological interest such as leaf tips, ligules, or inflorescence apices in the plant body.

**Figure 1.**
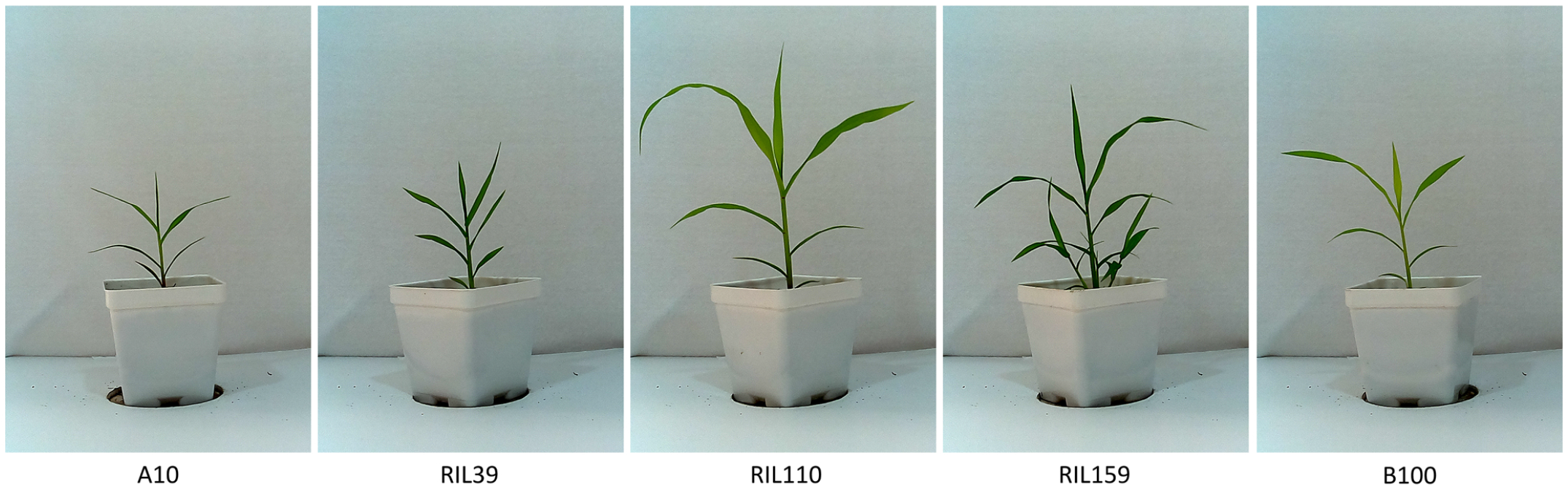
Example images of the genotypes that were used in this study at 15 days after germination.

### Generating Component Measurements from Pseudo-landmarks

Plant ontogenies are derived from various overlapping processes, with some describing variation at a component level, such as individual leaves or ligules along a stem, whereas others can be viewed as holistic attributes that govern the behavior of the axis in question, such as leaf number or flowering time. In order to model this variation, both the annotated pseudo-landmark identities and their x-y coordinate information were used for linear and nonlinear modeling approaches discussed at length below in the R language environment version 3.6.2: Dark and Stormy Night (R Core Team, 2019).

#### 1. Leaf Number and Flowering

Whole system measures such as leaf number and flowering time are the simplest measures to analyze and merely reflect counts based on the pseudo-landmark annotations. We used heading time, which is the appearance of an inflorescence in the flag leaf, as a proxy for flowering time, and treated it as a boolean value, where the state of heading begins as a zero value and shifts to one upon the day that the first inflorescence pseudo-landmark is observed. Leaf number focuses on identifying the highest numbered leaf annotation at each time point.

#### 2. Estimating Segmented Culm Heights from Ligule Landmarks

The first step in estimating height is to annotate the ligules corresponding to the main axis across developmental time, from which height can be calculated for the main stem as well as for the relative contribution of each phytomeric leaf sheath to total height. We use leaf sheath length to approximate internode length, as the internodes are not visible during the growth of the plant (Fig. 2). After ligule annotation, the central axis of the stem is defined using a linear regression of the ligule landmarks ranging from 1 to n with the formula:

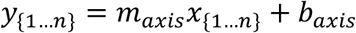

**Figure 2.**
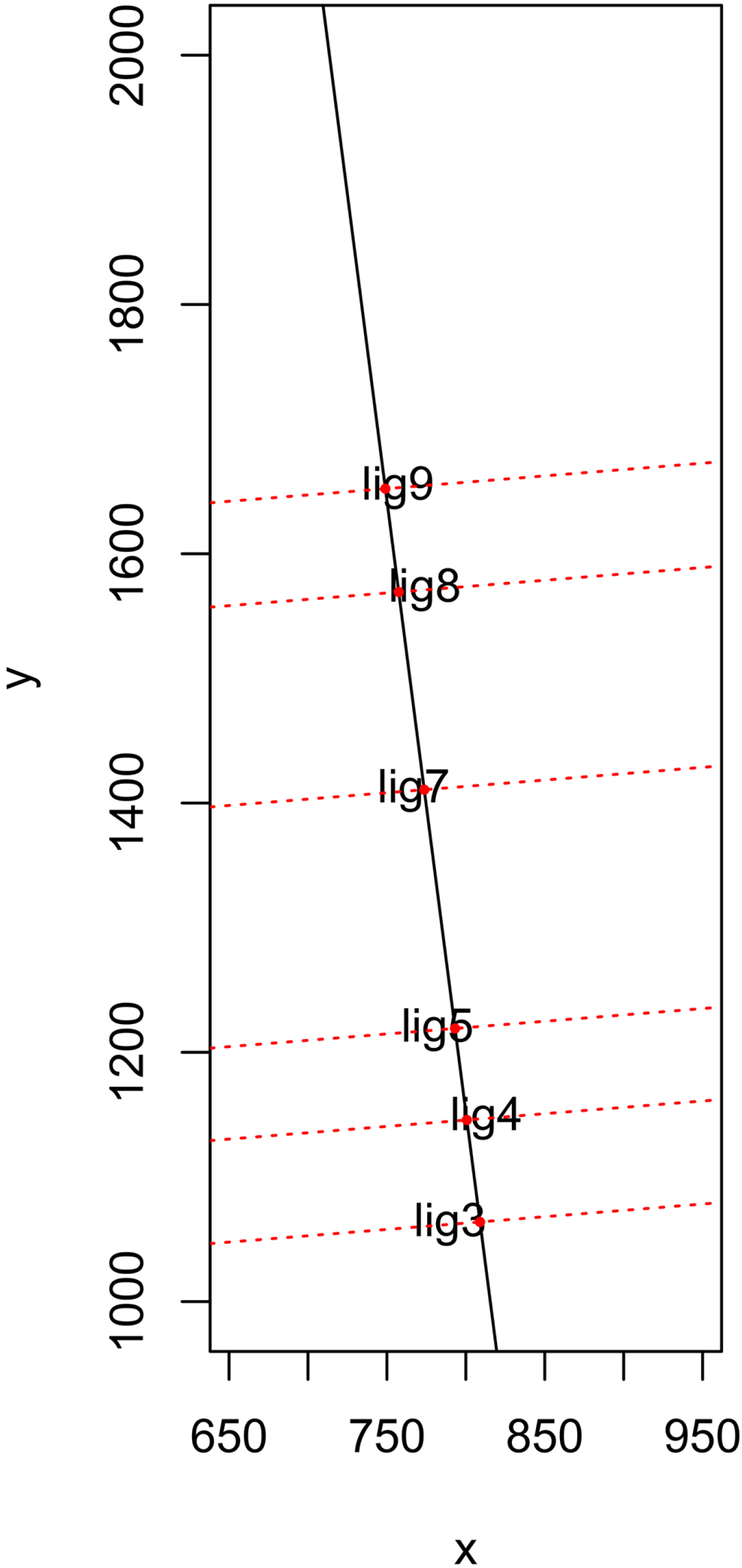
Example of the linear regression of a main axis derived from ligule pseudo-landmarks (black) as well as the subcomponent height segments at orthogonal angles along this regression (dotted red).

After estimating *m*_*axis*_ and *b*_*axis*_ the linear model can then be segmented into its subcomponents through the use of the original landmarks (*x, y*)_{1…;*n*}_. This is done by calculating the orthogonal axes to the previous linear model which contain the ligule landmarks (*x, y*)_{1…;*n*}_ *i*n which the orthogonal slope *m*_*orth*_ can be defined as:

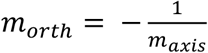

And the orthogonal intercept *b*_*orth*_ *c*an be defined as:

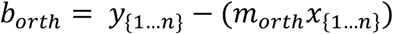

This results in an initial linear model that corresponds to the main culm itself as well as a range of orthogonal intersecting linear models which correspond to each ligule. From this, the vertices corresponding to these intersecting linear models can be computed following matrix multiplication of these corresponding slope and intercept values to produce vertices *v*_*{1…;n}*_. These vertices can then be used to estimate the relative contribution of each phytomer based on the distances between adjacent vertices, for using the formula:

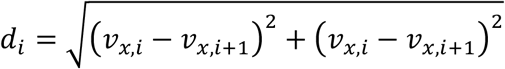

#### 3. Calculating Leaf Distances and Angles based on Ligule Vertices and Leaf Tip Positions

Following generation of the ligule vertices for measuring segmented culm height, leaf angles can be calculated with the corresponding leaf tip pseudo-landmark coordinates using the dot product method, which assumes the coordinates of all 3 vertices making up a triangle are known. In order to accommodate this assumption a 3^rd^ point, the apex, is arbitrarily generated de novo from the linear regression modeling the culm axis in which *x*_*apex*_ is solved for, given:

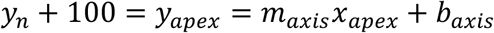

This result creates an additional point which will rest above the highest ligule (the elevation of which is represented by *y*_*n*_) allowing for this same apex coordinate to be used for all leaf angle measurements irrespective of the ligules location across the axis. The apex coordinates can then be used in conjunction with the ligule vertices and leaf tip coordinates given:

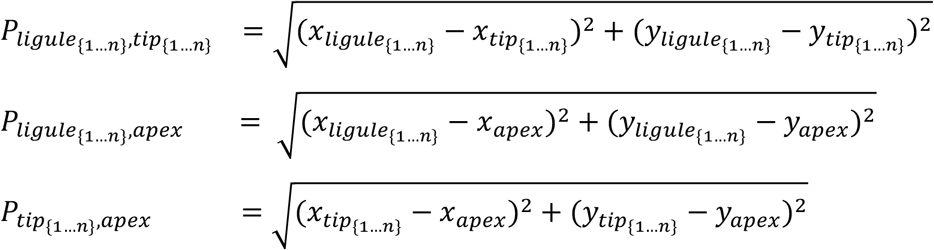

Which, together can then be used to solve for the leaf angle using the following dot product formula:

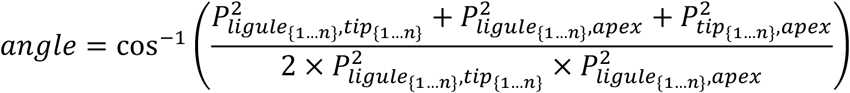

In the process of solving for leaf angle the leaf distance between the leaf tip and ligule is also solved for as *P*_*ligule*−*tip*_. Leaf distance is captured relatively poorly in this data set because of the window size of 1 cm used to assign unique pseudo-landmarks. This window size artificially generated a 2cm floor for all leaf distances measured.

### Function-Value Trait Modeling

The measurements generated from the pseudo-landmark data, were fitted with a sigmoidal growth model. This approach has the advantage of explaining factors such as the initiation of growth, magnitude of change, and attenuation of growth as parameters under a model instead of merely describing magnitudes of phenotypic change. To capture these interacting forces that propel and attenuate growth in a time-dependent manner, the following logistic formula was used:

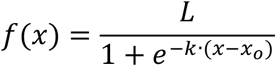

Although there are several variations on logistic equations the format of the equation above has the advantage of describing three parameters of interest, *L*, the plateau of the y-axis which describes attenuation of growth and the maximum size of an organ, *x*_*o*_, which describes the midpoint of growth when the rate of growth is the most rapid, as well as *k*, which defines the maximum rate of change at time *x*_*o*_. This model was chosen because it was structured in a manner to make it robust at describing both quantitative measures such as leaf numbers, leaf distances, leaf angles and segmented heights while still being able to also describe boolean measures such as heading date, when *L* plateaus at 1, without assuming unwarranted complexity. Parameter fitting for both whole shoot and individual structure FVT models was performed with the *nls* function in R with the port algorithm (R Core Team, 2019). The advantage provided by the use of port is that bounds for individual parameters can be specified to constrain possible fits and prevent convergence around a false optimum. In the case of whole shoot FVT modeling, the parameters *x*_*o*_ and *k* were allowed to remain relatively unconstrained while bounds were set for *L* based on the numbers of leaves or presence/absence of an inflorescence pseudo-landmark.

### Modeling of Individual Components

For individual modeling of components (i.e. leaves and ligules), parameter fitting entailed more nuanced constraints, given that interactions between these components are expected. The parameter *x*_*o*_ was constrained during parameter fitting so that each successive phytomer could not start within 2-3 days of the last (dependent on genotype). This constraint was based on the rate of phytomer deposition which had previously been calculated can be clearly seen in A10 (Fig. 3-B shows an example for accession A10, Supplementary Fig. 1 illustrates rates for all genotypes). Meanwhile, the parameter *L w*as constrained to not be larger than the observed measures. By contrast, parameter *k* was allowed to be unconstrained, and aids in maximizing the fit to the observed measures based on the constraints applied to *x*_*o*_ and *L*. An example for segmented height for genotype A10 is given in Figure 3.

**Figure 3.**
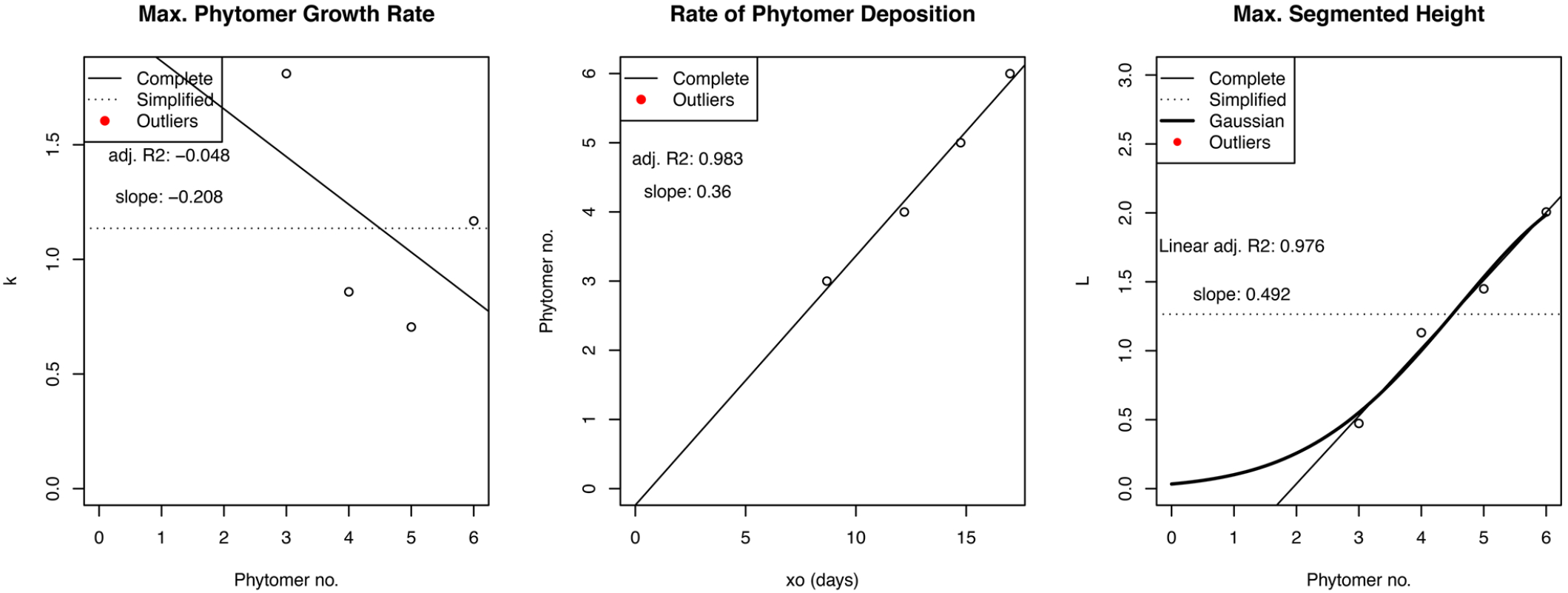
Example of the underlying structure of the FVT parameters *k, x*_*o*_, and *L* respectively underlying segmented height of individual models in genotype A10. Lines demonstrate various 2FVT approaches used in later trait modeling approaches. Mean value (dotted line), linear regression (solid line), gaussian model (boldened solid line).

### 2FVT Models

For models that correspond to consecutive structures such as segmented heights, leaf chord distances or leaf angles, second order trait modeling is performed using a variety of approaches for linear or nonlinear fitting across metamers (Table 1). The most intuitive of these approaches generates linear regressions of *L, x*_*o*_ and *k* relative to the structure’s phytomer position, which is hereafter referred to as the *complete linear* 2FVT method. These second-order linear models underlying the function-values for individual subcomponents are then used to predict the generalized 2FVT parameter at each phytomer position. By doing so, we hope to eliminate variation caused by measurement error etc. and obtain a more accurate measure of parameters for each phytomer. However, we found little correlation between *k* and phytomer position and thus also tested an alternative simplified model which used the mean *k* value for all phytomer positions which is hereafter referred to as the *simplified k* 2FVT method. The relative fits of the individual FVT models as well as the *complete linear* and *simplified k* 2FVT methods were compared. A linear regression used in both 2FVT methods was also not always adequate to explain variation in L, and two alternative strategies were performed. First, the linear regression of *L* was replaced with the mean *L* value and used in conjunction with the mean *k* value in order to minimize the parameter number (hereafter referred to as the *simplified k and L* 2FVT method). Second, a gaussian model that attempted to explain *L* by assuming more parameters,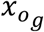, *k*_*g*_ *a*nd *L*_*g*_ *r*espectively, was also conducted using the following equation:

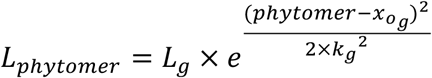

**Table 1.**

|  | ***k* parameters** | **$x_o$ parameters** | ***L* parameters** |
| --- | --- | --- | --- |
| ***simplified k and L*** | mean | slope, intercept | mean |
| ***simplified k*** | mean | slope, intercept | slope, intercept |
| ***complete linear*** | slope, intercept | slope, intercept | slope, intercept |
| ***simple gaussian*** | mean | slope, intercept | maxima, mean, standard deviation |

The gaussian equation provides a distribution around which starting and ending phenotypic measures can be fitted, in which *L*_*g*_ *i*s the highest measure realized, 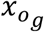 is the phytomer position around which *L*_*g*_ *i*s centered and *k*_*g*_ *i*s the standard deviation of this distribution. This gaussian derived *L* was then used alongside the mean *k* value for the *simplified k gaussian* 2FVT method (hereafter referred to as *simple gaussian* for brevity). The scripts used to parse pseudo-landmark data into informative measurements and to perform function-value trait modeling can be found on Github (https://github.com/jgerardhodge/2FVT).

### Comparison of 2FVT Models between Genotypes

Following the selection of an optimal 2FVT modeling strategy, comparisons between genotypes were able to be performed across their respective ontogenies. Agresti-Coull corrected Wald confidence intervals were estimated for each 2FVT model by genotype to enable comparisons against a standard curve, so that growth curves could be compared across the entirety of growth rather than merely at the inflection point of the curve. We first estimated a standard curve by calculating the central tendency between the parental genotypes through averaging their individual parameters underlying their 2FVT models. Second, we compared the confidence interval provided by the 2FVT model of each organ for each genotype and compared it to the standard curve of the central tendency between the parental genotypes. We simplified the outputs into a series of graphs which capture each attribute as well as by synthesizing their combined outputs into ‘in silico’ morphs. The scripts used to perform these analyses can be found at the previously mentioned Github repository.

### Accuracy of the 2FVT model

The accuracy of the 2FVT model for prediction was tested by comparing values for ligular plant height at flowering derived from the 2FVT models to independent measures of height measured in the original images using ImageJ. This was done by estimating the segmented heights for the phytomers based on the leaf number FVT model and summing these segmented heights to generate a 2FVT fitted culm height for each genotype. These 2FVT fitted estimations were then compared to the observed measures at flowering time using the *t*.*test* function in R, assuming a two-sided distribution of the observed measures for testing and that the fitted culm height represented the true mean (R Core Team, 2019).

### Ontogenetic Predictions from FVT and 2FVT Models for Flowering and Height

The curves derived from the FVT models were tested as a tool for predicting ontogeny, using flowering time predicted from when the leaf number FVT model plateaus as a demonstration. This prediction method was based on the observation that in four of the five genotypes (A10, RIL39, RIL110, RIL159) the flowering time observed was correlated with the asymptote of leaf number FVT models. Based on this insight, the first day at which the asymptote was reached and flowering was predicted to have occurred across these four genotypes was estimated using the first 70%, 75%, 80%, 85%, 90%, 95% as well as the complete time series dataset for each genotype. This approach was used to estimate the accuracy of predicting when the curve for leaf number should arrive at its asymptote given that this arrest of phytomer production is anticipated to correlate with flowering date and could be assessed based on the absolute difference of the observed and predicted values. B100 was not included as it had not flowered by the end of the experiment.

## Results

Leaf number and flowering time were modeled using a simple logistic regression, fitted to either count or boolean data respectively, from which leaf number and flowering *x*_*o*_, *k*, and *L* parameters were estimated (Fig 4). For one of the genotypes used in this study, B100, the flowering time and leaf number could not be captured by the pseudo-landmark dataset because the plants had not flowered before the end of the 30-day sampling window. Following the generation of trait measures from pseudo-landmark data, individual modeling of phytomers was performed for iterative growth attributes such as segmented height, chord length, leaf tip-ligule distances, and leaf angles. Although these fits could generally capture the trends of the data, they were often overly sensitive near the bounds of the sampling window for imaging when the full growth curve of the first and last organs could not be captured (Fig 5A). Due to this issue the second phytomer measures, although sampled in all cases, were omitted. A similar issue was seen near the end of the 30-day window, because recently initiated structures had not grown enough for their growth curves to have been captured sufficiently so that the underlying parameters could be estimated with confidence. In addition to these issues with parameter estimation for individual model fitted curves, this approach cannot predict organ growth rates for unsampled, or poorly sampled, organs, because no attempt is made to model underlying similarities between the curves. We developed the 2FVT nested regression strategy to take account of the relationship between the parameters underlying consecutive component models, in order to attempt prediction of under-sampled organs. In describing the individual parameters underlying each function-value trait as points along a curve which vary with respect to phytomer position, a holistic model can be developed that can extrapolate other components that either could not be observed or lacked reasonable resolution to accurately estimate, such as phytomer 2 (Fig 5B-D).

**Figure 4.**
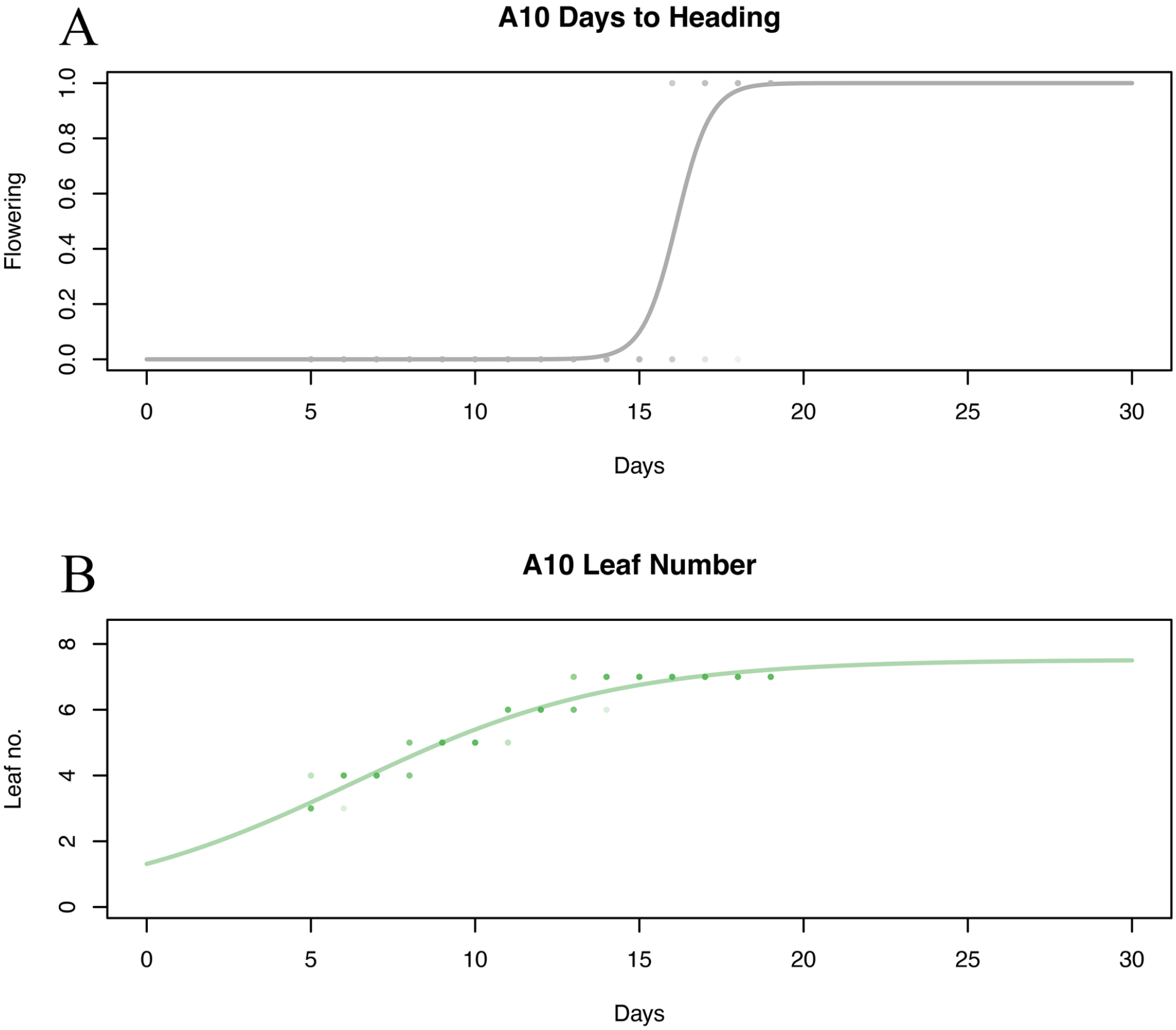
FVT model for the whole plant traits of (A) flowering and (B) leaf number in genotype A10. Points represent observed measures and line represents fitted model.

**Figure 5.**
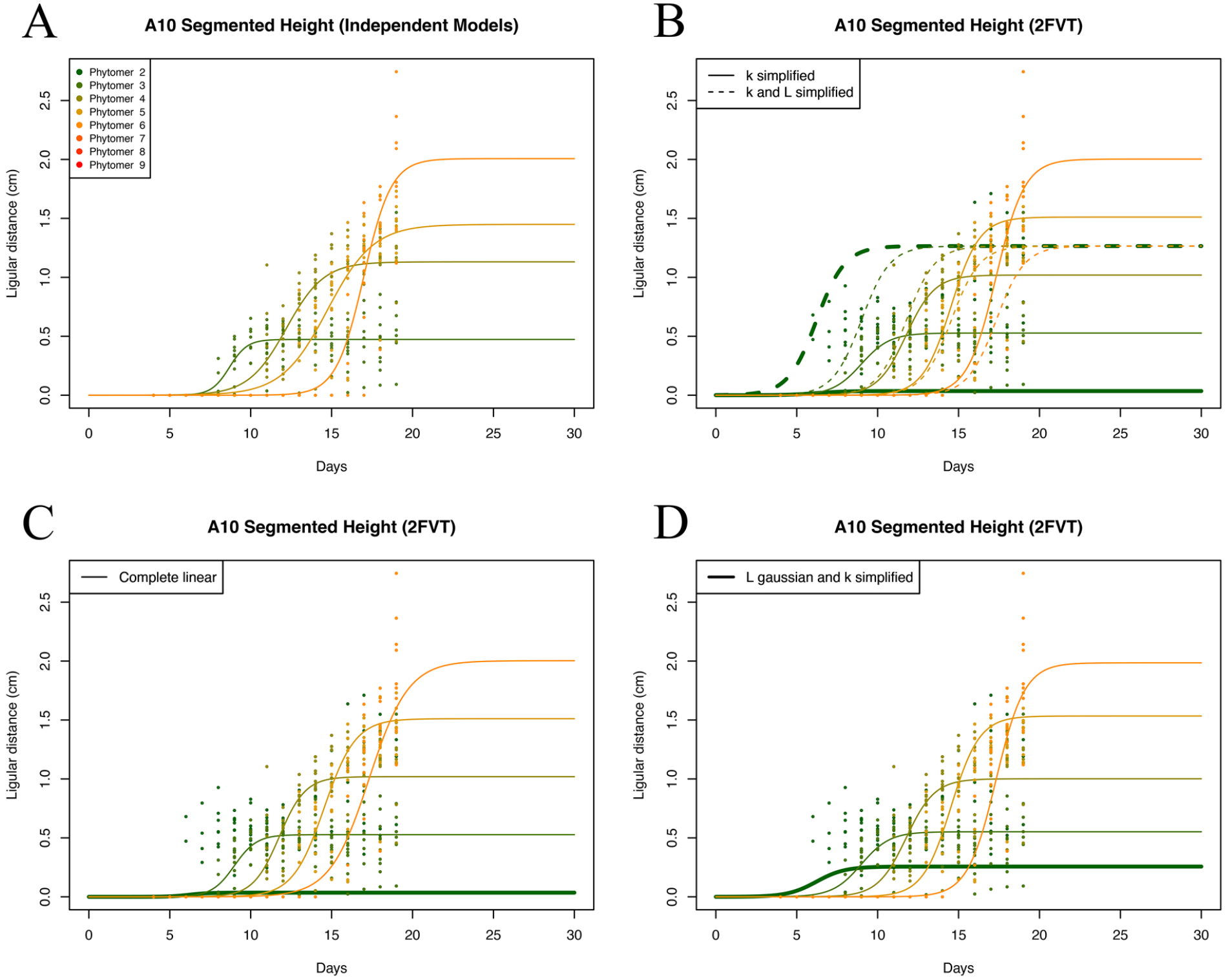
Segmented Height Individual (A) and 2FVT (B-D) modeling strategies for genotype A10. Points represent individual measures across replicates and lines the logistic regression fit with color gradient corresponding to phytomer position.

Various strategies were employed to estimate 2FVT model series that fell along a spectrum of complexity (Table 1). The simplest among these models, the *simplified k and L* method, assumed only 4 2FVT parameters, with one mean *k* and *L* value and a slope and intercept for the linear model for *x*_*o*_ (Fig. 5B). A somewhat more complex model, the *simplified k* method, assumed 5 2FVT parameters, with one mean *k* and a slope and intercept for *L* and *x*_*o*_, which were assumed to be linear (Fig. 5B). The *complete linear* strategy had 6 2FVT parameters, consisting of a slope and intercept all three parameters (Fig. 5C). An equally complex model was the *simple gaussian* model, which required 6 FVT parameters, 3 to capture the shape of *L* as a gaussian curve, 1 for a mean *k*, and 2 for the slope and intercept for *x*_*o*_ (Fig. 5D).

The accuracy of the various 2FVT approaches was estimated by both the discrepancy between values for traits provided by the models and independent measurements as well as by their predictions for phytomer models, where the values for that phytomer had been excluded. The first method revealed that the *complete linear* model was over-fitting on observed measures which left it susceptible to spurious predictions beyond this range. An artifact of this issue can be seen in Figure 5B-C where both the *complete linear* and *simplified k* grossly underpredict the size of the second phytomer. Other systemic errors were also observed in the *complete linear* 2FVT strategy such as inversion of growth processes (Fig 6). The *simplified k and L* model predicts an average phytomer size that does not fit the data well (Fig. 5B). However, the sim*ple gaussian* model appears able to both accurately represent the variation in final sizes of different structures when compared to their measures while also predicting sizes of structures such as the second phytomer which weren’t used to develop the 2FVT curves. Moreover, the *simple gaussian* model had an added advantage of not only specifying a growth ceiling that is reached at a given position but also a floor, which is absent in other models where *L* is assumed to be linear. This helps to prevent the erroneous predictions of negative growth in early phytomers (Fig. 3C).

**Figure 6.**
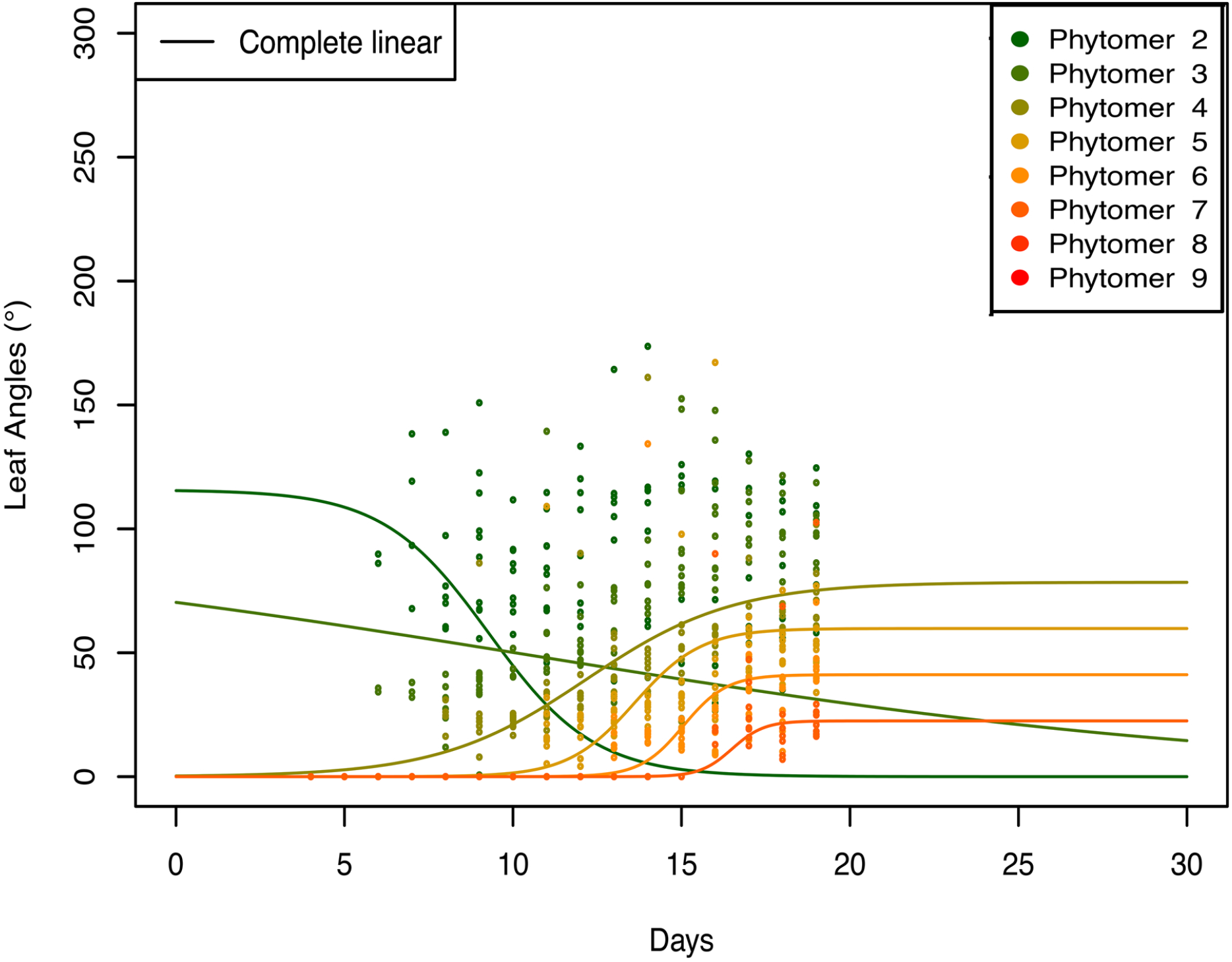
Demonstration of synergistic errors that can arise within the complete linear 2FVT model. Model inversion at phytomers 2 and 3 occurs due to interactions between the *k* and *L* linear fits.

To further test the accuracy of these varying 2FVT models, culm height was estimated using the *simple gaussian, complete linear*, and *simplified k* approaches by using the sum of the segmented heights at the terminal imaging day for each genotype. Observed ligular height measures were made across these terminal images for each of the genotype replicates in order to compare the distribution of the observed values against the 2FVT estimates (Figure 7). From this comparison the *simple gaussian* model was found to be a closer approximation to the observed values than the other 2FVT modeling approaches with a Student’s t-test finding no significant difference between the *simple gaussian* model and the observed measures in the majority of situations. In the cases where the *simple gaussian* model was estimated to differ significantly (RIL110 and B100) the absolute difference between the *simple gaussian* and observed mean heights was less than 2 cm which was generally an order of magnitude less than the other 2FVT modeling approaches (Supplementary Table 1). Based on the poor predictive performance of the *complete linear* and *simplified k* 2FVT models for the beginning and end elements of the phytomer series (Figs. 5B-C & 6) as well as the high accuracy of the *simple gaussian* model in approximating the observed measures for culm height (Fig. 7) the *simple gaussian* model was determined to be the most accurate for trait modeling in this system.

**Figure 7.**
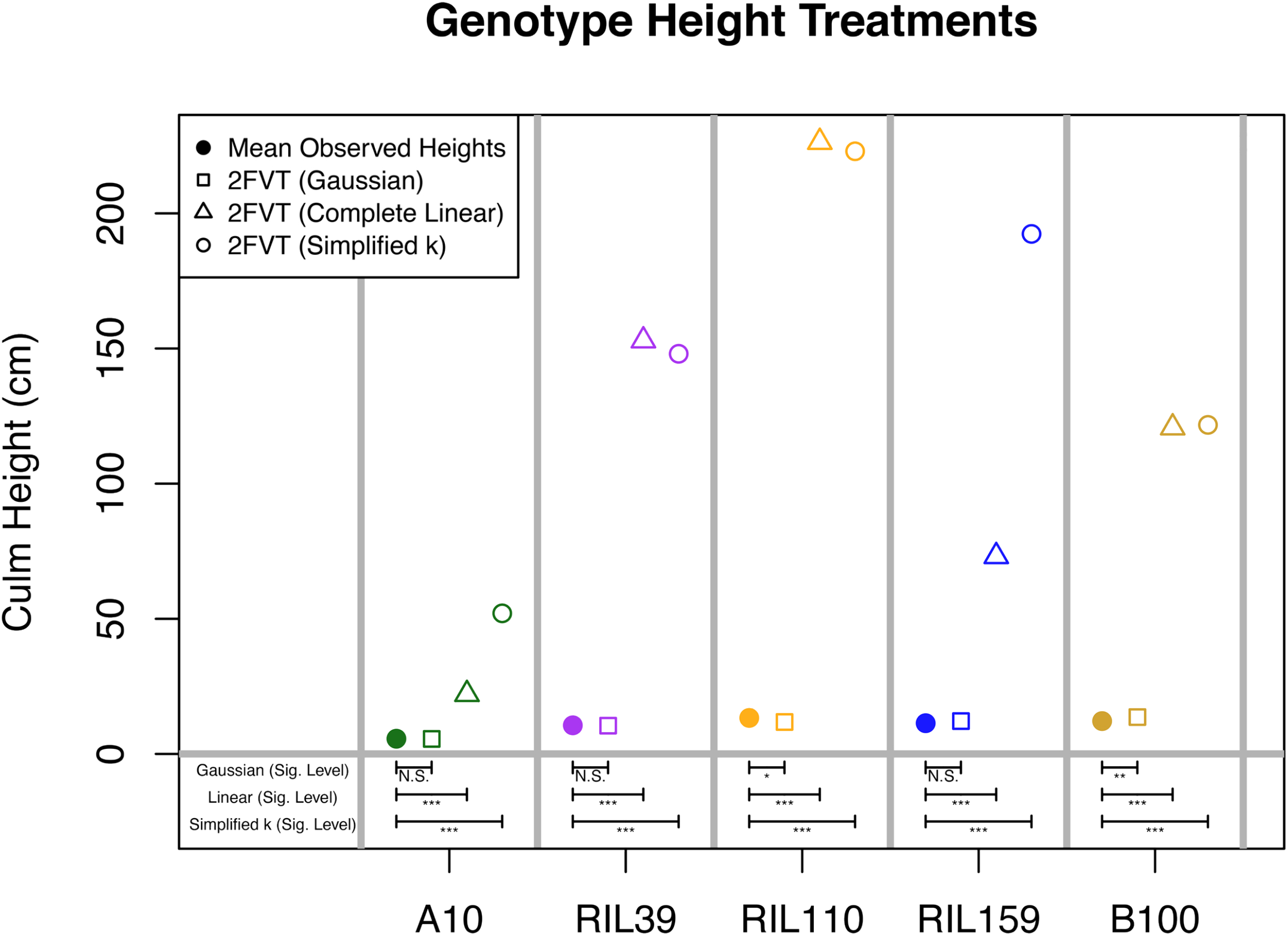
Accuracy of 2FVT modeling approaches at termination of time course imaging for each genotype. The mean value of biological replicates (solid circle) either at flowering (A10, RIL39, RIL110, RIL159) or at the end of the growing experiment (B100) were compared to the culm heights predicted by the simple gaussian (square), complete linear (triangle), or simplified k (hollow circle) 2FVT models based on the sum of the segmented heights at this time point. Statistical comparisons between 2FVT predicted means and the observed measures were made by performing T-test’s the significance levels of which are denoted below with brackets indicating each comparison.

We can compare individual parameters from the *simple gaussian* 2FVT models for the different genotypes in order to understand how morphological variation may be generated (Supplementary Fig. 1, Supplementary Table 2). Prior to modeling growth of individual structures, genotype specific signatures can already be seen in these 2FVT parameters as outlier trends (Supplementary Fig. 1). For example, RIL110 can generally be seen to have a steeper *x*_*o*_ slope, particularly in the segmented height trait space when compared to all other genotypes, which corresponds to higher *L* values at equivalent phytomer positions in this background. In addition, the L values estimated for leaf angle in RIL39 can be seen to be relatively flat compared to other genotypes which corresponds to a shallower *x*_*o*_ slope in this trait space. Based on the evidence seen in these parameter models it seems that phenotypic variation between genotypes can already been seen as underlying the parameters of the growth models themselves. Moreover, there is an added power to these parameters in that they allow for fitting intermediate states between multiple genotypes through averaging them allowing for smooth curves to compare against as standards. This is demonstrated by the transgressive mean values which represent that averaged A10 and B100 parameters (Supplementary Fig. 1). As demonstrated by the 2FVT estimated curve for leaf number, the transgression test standard curve generated by using the individual transgressive means of the parental parameters can readily produce a smoothed mean phenotypic estimation (Fig. 8). This smoothed curve is able to accurately track the general tendency of the parental lines over time which can then be compared to the confidence intervals of individual genotypes (Fig. 8). Moreover, this approach works both at the scale of whole shoot level processes such as leaf number and flowering and at the level of individual organs or structures of interest (See Supplementary Data 1).

**Figure 8.**
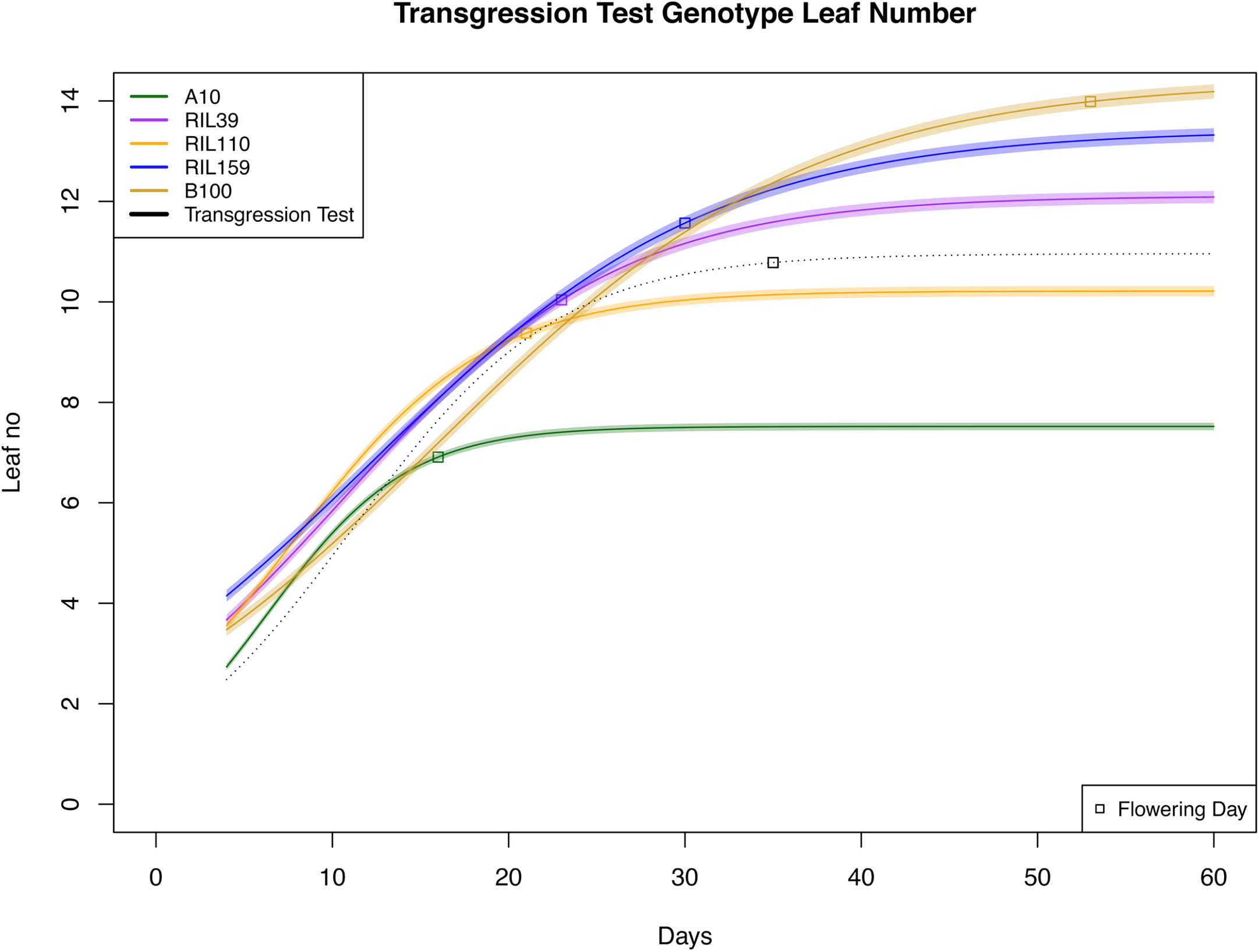
Example of the estimated phenotypic values for leaf number derived from the FVT parameters alongside 95% Wald confidence intervals. Genotypes are represented by color and the transgressive standard representing the mean values of A10 and B100 is denoted as a dotted black line. Mean fitted flowering dates are also shown for each genotype beneath the estimated growth curves.

The 2FVT parameter estimates clearly represent a multidimensional dataset that is difficult to interpret collectively when it is depicted in a manner that removes the underlying morphology that interconnects these trait spaces (Supplementary Fig. 1). With this issue in mind, efforts were made to simplify these outputs as ‘in silico’ morphs to compare growth attributes such as chord leaf distance, leaf angle, height, and flowering simultaneously (Fig. 9, Supplementary Movie). This approach has the advantage of being able to describe not only how genotypes vary in general morphology but also can describe the manner in which growth varies across the body plan to achieve it and when. An example is given for plant height, as this is a common trait that is often studied because of its high heritability and correlation with flowering. At day 15 the lines A10, RIL39, RIL159 and B100 are all roughly equivalent to one another in maximum height but vary in the number of height segments, with A10 and B100 having one fewer phytomer than the two RILs. At day 25 differences in height segment characteristics become even more pronounced when RIL39, RIL110 and B100 are equivalent in height despite the fact that RIL110 achieves this variation by producing fewer, longer height segments in contrast to RIL39 which has more numerous, short segments. Which height segments are elongated compared to others along the axis also varies between genotypes. In comparison to the RILs, B100 produces a fairly homogeneous array of moderately longer height segments and will ultimately overtake them in height, because the RILs have transitioned to flowering while B100 is still vegetative and undergoing internode expansion. In this way the 2FVT model allows for an appreciation of the complexity underlying a trait such as height.

**Figure 9.**
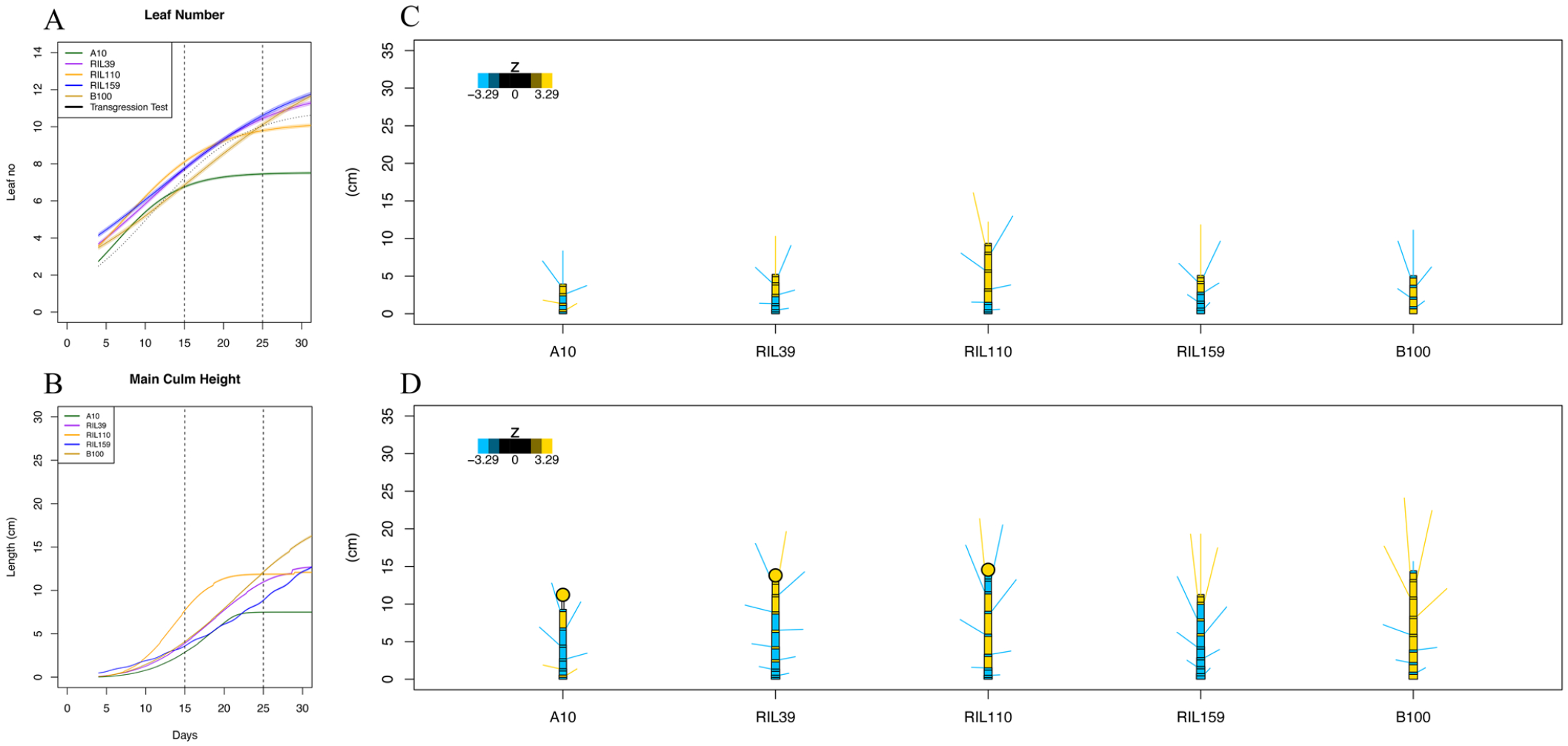
Whole plant traits of leaf number and plant height to compare against in silico morphs at the denoted time slices (dotted lines). (A) Leaf number over time with genotypes denoted by color. (B) Plant height with genotypes denoted by color. (C-D) In silico morphs demonstrating the variety of 2FVT phenotypes simultaneously by genotype at days 15 and 25 respectively. Coloring of in silico structures denotes underperformance (blue) or overperformance (yellow) with respect to the transgressive standard curve. Scoring colors for these performance statistics can be interpreted as follows: ligules coloring represents leaf angles, inter-ligular segments represent segmented height, blade color represents leaf distances, inflorescence circle represents flowering.

In addition to the advantages of being able to more accurately model growth, the 2FVT model can also be used to predict phenotypes estimated from the parameters. We tested the predictive ability of the models for flowering time using a strategy of removal of 0-30% of the terminal time series data for leaf counts (Table 2). Generally, the predicted heading dates fell within at least a week of the observed value, with many even falling much closer when the difference between the observed and predicted values are compared. Using the models to predict flowering time in B100, which had not flowered by the end of the experiment, resulted in a prediction of flowering of flowering approximately 53 days after planting with approximately 16 leaves being produced (Fig. 8), which our previous experience with this genotype suggested was reasonable.

**Table 2.**

|  | **A10** | **RIL39** | **RIL110** | **RIL159** | **B100** |
| --- | --- | --- | --- | --- | --- |
| **Observed Flowering** | **16.1** | **22.6** | **20.9** | **29.5** |  |
| 70% Prediction | 20.8 | 28.1 | 24 | 27.9 | - |
| $\vert \Delta\vert $ | 4.7 | 5.5 | 3.1 | 1.6 | - |
| 75% Prediction | 24 | 28.1 | 17.6 | 25 | - |
| $\vert \Delta\vert $ | 7.9 | 5.5 | 3.3 | 4.5 | - |
| 80% Prediction | 24 | 26.9 | 22.3 | 24.8 | - |
| $\vert \Delta\vert $ | 7.9 | 4.3 | 1.4 | 4.7 | - |
| 85% Prediction | 21.9 | 27.4 | 26.9 | 25 | - |
| $\vert \Delta\vert $ | 5.8 | 4.8 | 6 | 4.5 | - |
| 90% Prediction | 24.7 | 27.6 | 20.6 | 25 | - |
| $\vert \Delta\vert $ | 8.6 | 5 | 0.3 | 4.5 | - |
| 95% Prediction | 21.1 | 28.3 | 29.3 | 29.3 | - |
| $\vert \Delta\vert $ | 5 | 5.7 | 8.4 | 0.2 | - |
| 100% Prediction | 17.7 | 24.4 | 20.6 | 29.5 | 52.5 |
| $\vert \Delta\vert $ | 1.6 | 1.8 | 0.3 | 0 | - |

## Discussion

The 2FVT method is based on a relatively simple assumption: that there is one single underlying growth model governing the developmental models of sequential organs of interest and that the individual models explaining the growth and development of each member of an ontogenetic series, such as a particular leaf or segmented height metamer, are simply the derivatives of the 2FVT model.

Several 2FVT models were used to analyze sequentially produced structures, but only one, the *simple gaussian* model, provided an adequate fit. There are various reasons for the failures of the other three models. The *simplified k and L* method erroneously assumes that all structures at all positions are homogeneous and, although this makes for a much easier model to parameterize, it ignores known patterns in plant morphology such as ontogenetic drift and heteroblasty (Evans 1972, Steeves & Sussex, 1989). The *simplified k* method also errs in over-generalizing the pattern of *L* by presuming it to be indefinitely linear. This leads to both under estimating organ size in early phytomers and over estimating size in higher order positions along an axis (Supplementary Data 1). The *complete linear* model also suffers from the same artifacts of over and under estimations of size from assuming *L* is linear but is further affected by *k* being fitted as a linear model as well. This can result in fallacious model at the boundary phytomer positions where the values of *k* and *L* are the most extreme (Fig. 6). By contrast, the *simple gaussian* model provides both a floor and ceiling for maximum sizes which effectively serve as guard rails and help to produce measures that more realistically approximate observed measures of growth. It should also be noted that the capacity for successive organs to be fitted with a single exponential parameter (i.e. simplifying *k* to a mean value that’s shared across phytomers) has previously been used successfully for modeling exponential growth within stem elongation (Fournier & Andrieu 2000).

The use of a gaussian model as a descriptor of *L* probably best fits the growth strategy of a determinate plant, where the shoot apical meristem is prioritized for reproduction during phase change, and is unlikely to be generalizable across all land plants. One might suspect on their face that plants such as *Vitis* or *Passiflora*, which exhibit indeterminate growth (Chitwood et al. 2015, Chitwood et al. 2017), might be better described with an asymptotic model for L, given the fact that environmental pressures rather than hard set ontogenic constraints on phytomer number are usually what lead to the cessation of growth. However, previous studies which surveyed the the a-seasonal internode expansion in the branches of Hevea over a growing season showed that that a gaussian pattern could still be recovered so the appropriate application of this model in perennial indeterminate systems remains an open question (De Reffye & Kang 2011).

Heteroblasty may in fact also offer further complexity in which different phases of growth require distinct, phase-dependent gaussian distributions as could be easily envisioned by the morphological extremes seen in nature such as the clonal branching habits of tank bromeliads in which lateral branches exhibit notably longer internodes as they initially grow out compared to the rosette tanks which exhibit hardly any internode expansion thereafter (Sampaio et al. 2004). The gaussian model underlying *L* in this study is likely correlated with life history strategies in annuals, given that they have organ ontogenies that evade suboptimal growth conditions by rapidly completely their life cycle in a finite window when the environment is optimal (Costa et al. 2012, Vanhaeren et al. 2015). Such models could well serve as a basis for exploring fitness landscapes with respect to vegetative morphotypes and explain how strategies are optimized within genetic mapping populations or across species boundaries.

As shown by the genotype comparisons, the 2FVT model identifies not only by how much a genotype’s traits exceed some expectation, but also when this occurs. The transgressive segregation test explicitly asks when and where each genotype’s phenotypic values exceed that of the mean tendency between the parents, and shows marked variation between genotypes. The in silico models presented in Figure 9 combine traits that likely have at least partially independent ontogenetic variation and show how these co-vary during growth, while also allowing comparisons between genotypes (allomorphosis – Niklas 1994). Although modeling genotypes through time could clearly be expected to correspond to specific allometric comparisons at either a particular stage or orthologous organ the inclusion of time provides insight as to when traits of specific genotypes become outliers, but more importantly provide a context as to how this is achieved. As seen in the in-silico morphs at day 25 (Fig. 9D) three genotypes (RIL39, RIL110, B100) achieve equivalent heights at similar points in their respective ontogenies although the degree to each individual segmented height phytomer and their overall number varies which may not only have impacts on their desired stature but also provide differences in their overall rigidity. However, the similarities in height are transient, because B100 has not completed its growth at this stage, whereas RIL39 and RIL110 are nearing the end of their vertical growth. In studies focused on allometric comparisons that are made at only a few time points, this variation would be lost, and therefore the 2FVT approach provides a key tool for modeling phenotypes through time both to explore comparative morphology and dissect the genetic basis of individual ontogenetic progressions.

The predictive power of taking ontogenetic series into account may be indicative of the fact that these function-value traits, be they whole plant or second-order models, have some inherent “biological meaning” (Niklas 1994). Given that the 2FVT approach relies heavily on phytomer-based approximations it seems reasonable to hypothesize that these parameters are linked to ontogenetic decisions of primordia initiated along the shoot apical meristem. There is some evidence already from other function-value trait studies that these parameters are heritable (Baker et al. 2015). Genetics studies have also proven informative in dissecting the functional mechanisms of heteroblastic series and allometric shifts between taxa and would provide an obvious prospect for mining the 2FVT derived allomorphic variation within a mapping population (Bongard-Pierce et al. 1996, Langlade et al. 2005, Demmings et al. 2019). The parameters of the 2FVT models are also affected by environment and these values could be anticipated to change in response to environmental perturbations thus providing another fruitful avenue for consideration, in which the gene-by-environment interactions are rooted in changes in ontogeny (Doust et al. 2014). Both of these subjects will pose a fruitful area for future research in which the scale of this modeling technique is expanded in order to better ascertain both the mechanistic and genetic basis of these parameters in their association with ontogeny. With the ease of generating high resolution data, it will become increasingly easy to incorporate ontogenetic shifts into models of organ growth. The possibility of synergies between 2FVT trait spaces also remains a subject of interest.

## Acknowledgements

We would like to William M. Hammond for his insightful feedback at various stages of this manuscript.

---

**Supplementary Figure 1**. Example of 2FVT parameter fits across phytomers with genotypes represented by color. Transgressive mean values of parameters for the A10 and B100 parental genotypes shown as a dotted black line.

**Supplementary Movie 1**. Animation demonstrating the 2FVT derived measures through the ontogenies of the genotypes used when tested against the standard curve of the transgressive mean of the parental (A10 & B100) genotypes. Black are approximating the mean at each point in time whereas blue are excessively underperforming compared to the standard curve and yellow are excessively overperforming.

## Notes

### Competing Interest Statement

The authors have declared no competing interest.

## References

Arber A. (1946). Goethe’s Botany. Chronica Botanica 10(2): 67–124.

Baker RL, Leong WF, Brock MT, Markelz RJC, Covington MF, Devisetty UK, Edwards C, Maloof J, Welch S, Weinig C. Modeling development and quantitative trait mapping reveal independent genetic modules for leaf size and shape. New Phytol. 208: 257–268.

Bongard-Pierce DK, Evans MS, Poethig RS. (1996). Heteroblastic features of leaf anatomy in maize and their genetic regulation. Intd. J. Plant Sci. 157(4): 331–40.

Chitwood, DH, Klein LL, O’Hanlon R, Chacko S, Greg M, Kitchen C, Miller AJ, Londo JP. (2015). Latent developmental and evolutionary shapes embedded within the grapevine leaf. New Phytol. 210: 343–55. doi: https://doi.org/10.1111/nph.13754

Chitwood, D. & Otoni WC. (2017). Divergent leaf shapes among Passiflora species arise from a shared juvenile morphology. Plant Direct 1(5). doi: https://doi.org/10.1101/067520

Costa MMR, Yang S, Critchley J, Feng X, Wilson Y, Langlade N, Copsey L, Hudson A. (2012). The genetic basis for natural variation in heteroblasty in Antirrhinum. New Phytol. 196(4): 1251–59.

De Reffye P & Kang M. (2012). Stochastic modeling of tree annual shoot dynamics. Ann. Forest Sci. 69: 153–65.

Demmings EM, Williams BR, Lee C, Barba P, Yang S, Hwang C, Reisch BI, Chitwood DH, Londo JP. (2019). Quantitative trait locus analysis of leaf morphology indicates conserved shape loci in grapevine. Front Plant Sci. 10:1373 doi: https://doi.org/10.3389/fpls.2019.01373

Doust AN, Lukens L, Olsen KM, Mauro-Herrera M, Meyer A, Rogers K. (2014). Beyond the single gene: how epistasis and gene-by-environment effects influence crop domestication. PNAS 111(17): 6178–83.

Evans GC. (1972). The quantitative analysis of plant growth. Berkley and Los Angelas, USA. University of California Press.

Feldman MJ, Paul RE, Banan D, Barrett JF, Sebastian J, Yee M, Jiang H, Lipka AE, Brutnell TP, Dinenny JR, Leake ADB, Baxter I (2017). Time dependent genetic analysis links field and controlled environment phenotypes in the model C4 grass, Setaria. PloS Genet. 13(6): e1006841. doi: 10.1371/journal.pgen.1006841

Fournier C & Andrieu B. (2000). Dynamics of the elongation of internodes in maize (Zea mays L.): analysis of phases of elongation and their relationships to phytomer development. Ann. Bot. 86: 551–63.

Hammond D. (1941). The expression of genes for leaf shape in Gossypium hirsutum L. and Gossypium arboreum L. I. The expression of genes for leaf shape in Gossypium hirsutum L. Amer. J. Bot. 28(2): 124–38.

Hodge JG, Li Q, Doust AN. (preprint). De novo homology assessment from landmark data: A workflow to identify and track segmented structures in plant time series images. doi: https://doi.org/10.1101/2021.02.21.432162

Huxley JS, Needham J, Lerner IM. (1941). Terminology of relative growth rates. Nature 146: 618.

Jones CS. (1999) An essay on juvenility, phase change, and heteroblasty in seed plants. Int. J. Plant Sci. 160(S6): S105–11.

Langlade NB, Feng X, Dransfield T, Copsey L, Hanna AI, Thébaud C, Bangham A, Hudson A, Coen E. (2005). Evolution through genetically controlled allometry space. PNAS 102(29): 10221–26.

Mauro-Herrera M & Doust AN (2016). Development and Genetic Control of Plant Architecture and Biomass in the Panicoid Grass, Setaria. PloS one 11(3): e0151346. doi: 10.1371/journal.pone.0151346

McCormick RF, Truong SK, Mullet JE. (2016). 3D Sorghum reconstructions from depth images identify QTL regulating shoot architecture. Plant Physiol. 172: 823–34.

Nijhout HF & German RZ. (2012). Developmental causes of allometry: new models and implications for phenotypic plasticity and evolution. Integr. Comp. Biol. 52(1): 43–52.

Niklas KJ. (1994). Plant allometry: the scaling of form and process. Chicago, USA: University of Chicago Press.

Miao C, Pages A., Xu Z, Rodene E, Yang J, & Schnable JC (2020). Semantic segmentation of Sorghum using hyperspectral data identifies genetic associations. Plant Phenomics 4216373: 1–12.

Pound MP, Atkinson JA, Townsend AJ, Wilson MH, Griffiths M, Jackson AS, Bulat A, Tzimiropoulus G,Wells DM, Murchie EH, Pridmore TP, French AP (2017). Deep machine learning provides state-of-the-art performance in image-based plant phenotyping. Gigascience 6(10): gix083. doi: 10.1093/gigascience/gix083

Prusinkiewicz P, Erasmus Y, Lane B, Harder LD, Coen E. (2007). Evolution and development of inflorescence architectures. Science 316: 1452–56.

R Core Team (2019). R: A language and environment for statistical computing. Vienna, Austria: R Foundation for Statistical Computing. https://www.R-project.org/

Sampaio MC, Araújo TF, Scarano FR, Stuefer JF. (2004). Directional growth of a clonal bromeliad species in response to spatial habitat heterogeneity. Evol. Ecol. 18: 429–42.

Sinnott EW. (1921). The relation between body size and organ size in plants. Am. Nat. 55(640): 385–403.

Steeves TA & Sussex IM. (1989). Patterns in plant development. New York, USA: Cambridge University Press.

Vanhaeren H, Gonzalez N, Inzé D. (2015). A journey through a leaf: phenomics analysis of leaf growth in Arabidopsis thaliana. Arabidopsis Book 13: e0181. doi: https://doi.org/10.1199/tab.0181/

Zotz G, Wilhelm K, Becker A. (2011). Heteroblasty-a review. Bot. Rev. 77: 109–51. doi: https://doi.org/10.1007/s12229-010-0962-8

