## Supplementary Data 1 for "Wheels within wheels, Nested Function-value Traits as a Tool for Modeling Ontogeny"

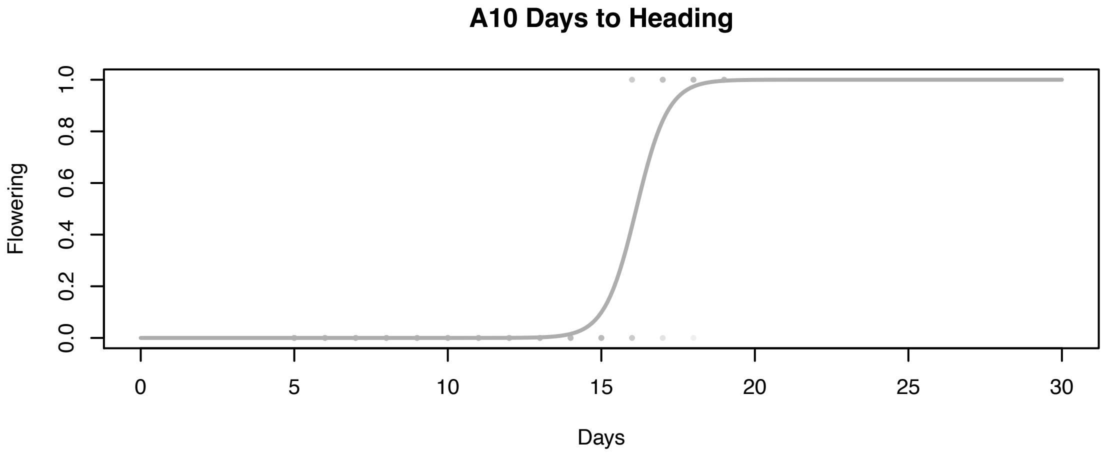
A10 Days to Heading
Flowering
0.0
0.2
0.4
0.6
0.8
1.0
0
5
10
15
20
25
30
Days

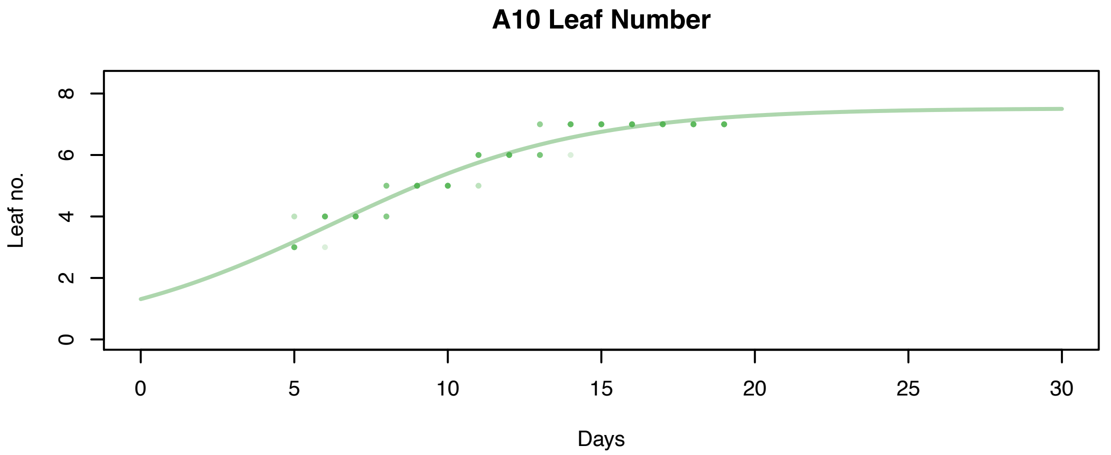
A10 Leaf Number
Leaf no.
0
2
4
6
8
0
5
10
15
20
25
30
Days

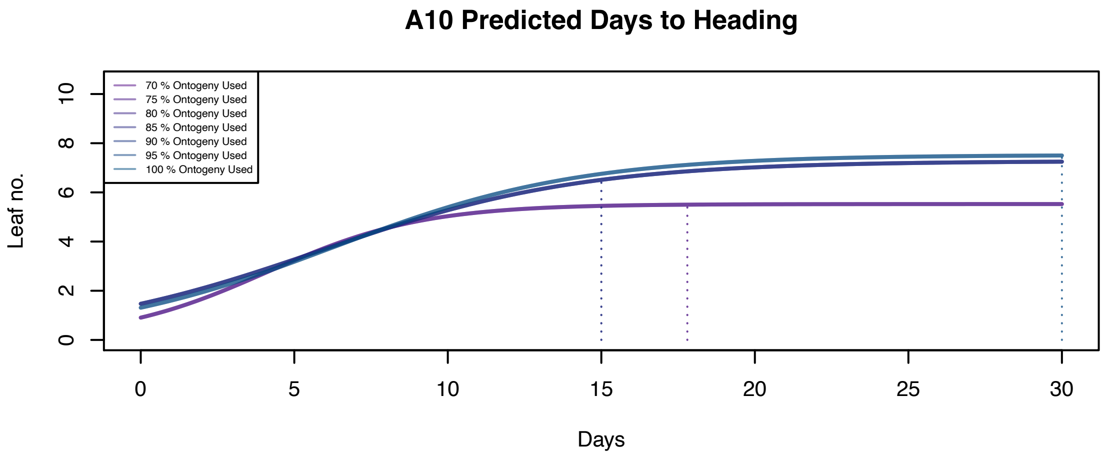
A10 Predicted Days to Heading
Leaf no.
70 % Ontogeny Used
75 % Ontogeny Used
80 % Ontogeny Used
85 % Ontogeny Used
90 % Ontogeny Used
95 % Ontogeny Used
100 % Ontogeny Used
0
2
4
6
8
10
0
5
10
15
20
25
30
Days

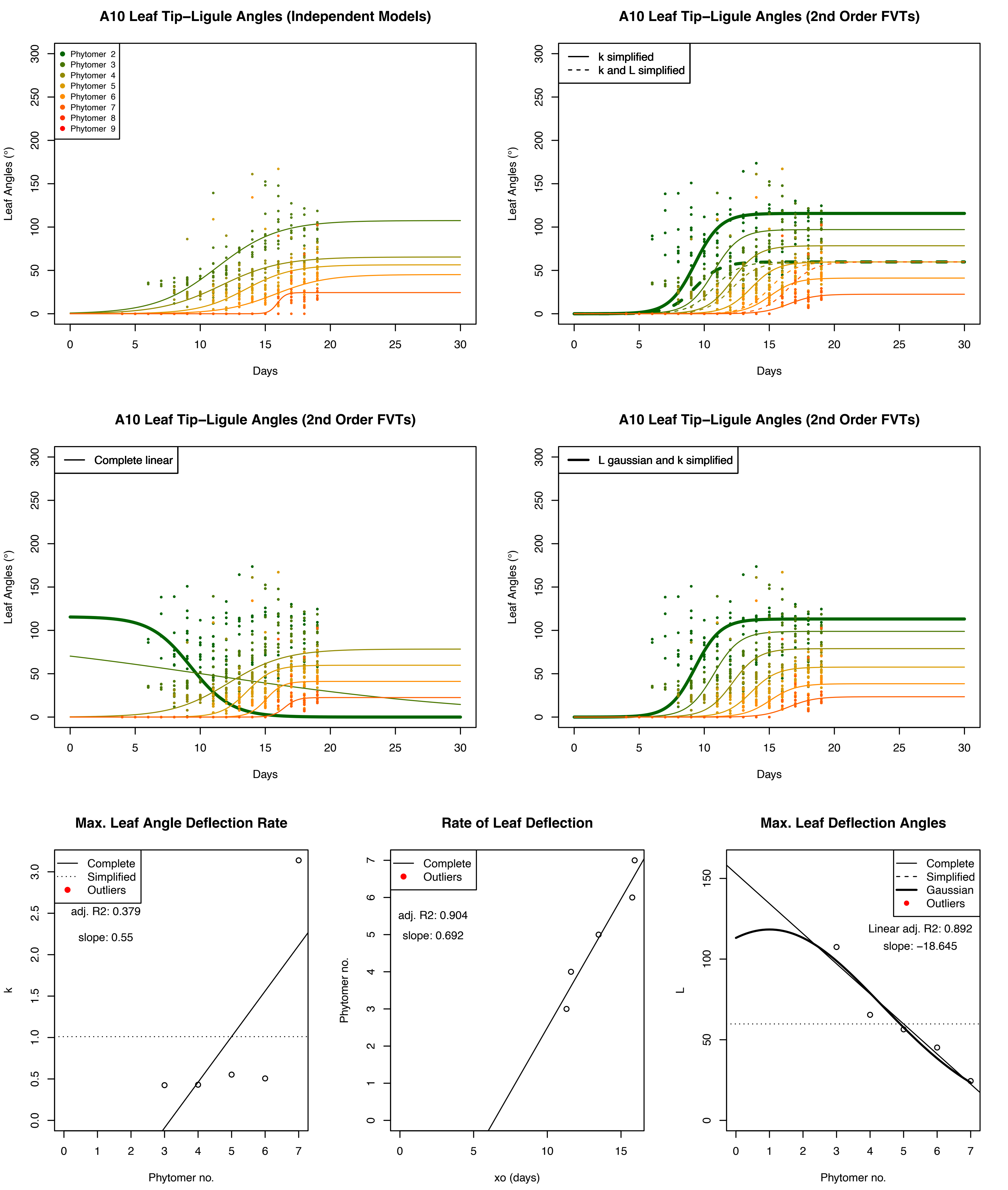
A10 Leaf Tip–Ligule Angles (Independent Models)
Phytomer 2
Phytomer 3
Phytomer 4
Phytomer 5
Phytomer 6
Phytomer 7
Phytomer 8
Phytomer 9
Leaf Angles (°)
Days
A10 Leaf Tip–Ligule Angles (2nd Order FVTs)
k simplified
k and L simplified
Leaf Angles (°)
Days
A10 Leaf Tip–Ligule Angles (2nd Order FVTs)
Complete linear
Leaf Angles (°)
Days
A10 Leaf Tip–Ligule Angles (2nd Order FVTs)
L gaussian and k simplified
Leaf Angles (°)
Days
Max. Leaf Angle Deflection Rate
Complete
Simplified
Outliers
adj. R2: 0.379
slope: 0.55
k
Phytomer no.
Rate of Leaf Deflection
Complete
Outliers
adj. R2: 0.904
slope: 0.692
Phytomer no.
xo (days)
Max. Leaf Deflection Angles
Complete
Simplified
Gaussian
Outliers
Linear adj. R2: 0.892
slope: −18.645
L
Phytomer no.

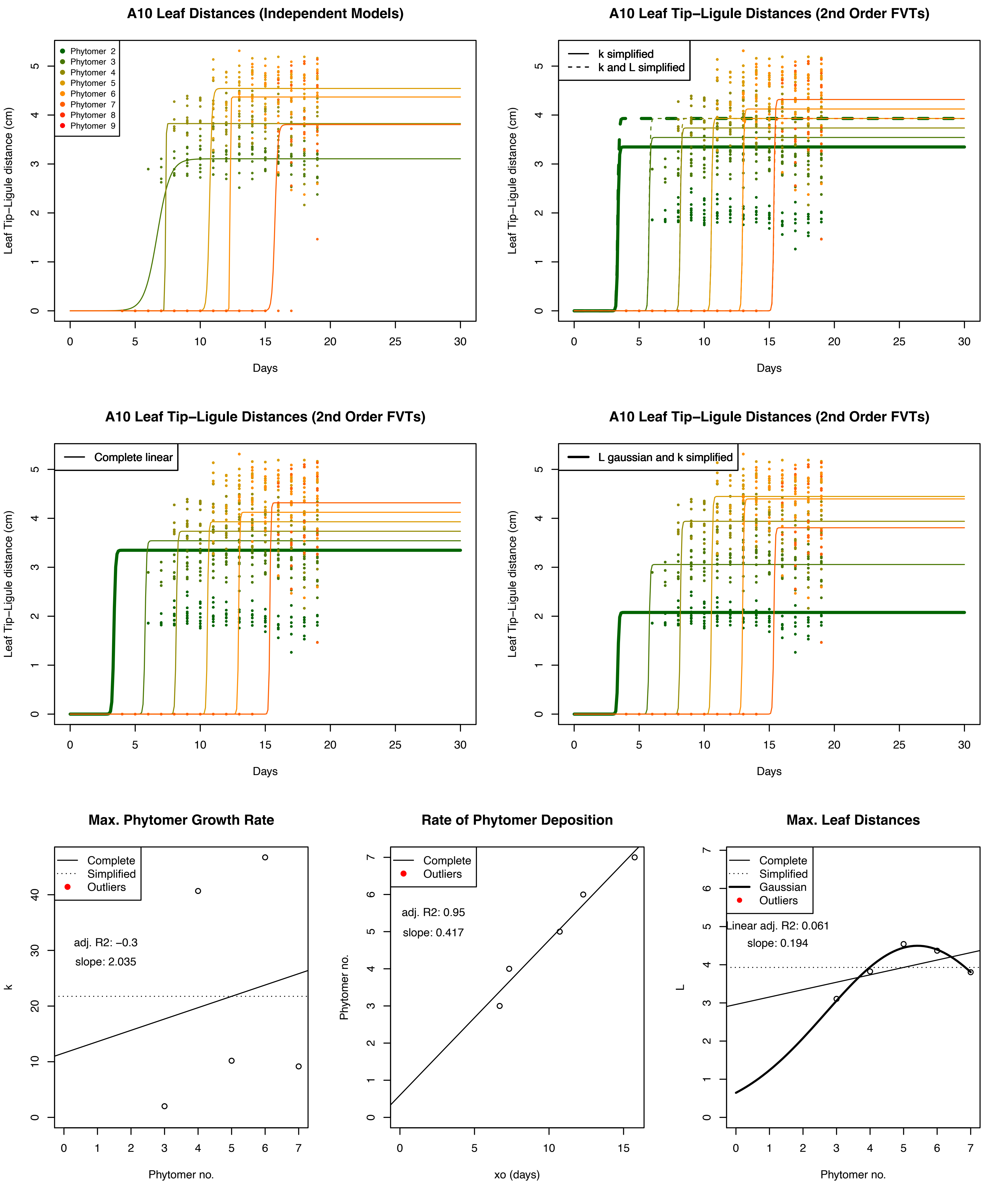
A10 Leaf Distances (Independent Models)
Phytomer 2
Phytomer 3
Phytomer 4
Phytomer 5
Phytomer 6
Phytomer 7
Phytomer 8
Phytomer 9
Leaf Tip−Ligule distance (cm)
Days
A10 Leaf Tip−Ligule Distances (2nd Order FVTs)
k simplified
k and L simplified
Leaf Tip−Ligule distance (cm)
Days
A10 Leaf Tip−Ligule Distances (2nd Order FVTs)
Complete linear
Leaf Tip−Ligule distance (cm)
Days
A10 Leaf Tip−Ligule Distances (2nd Order FVTs)
L gaussian and k simplified
Leaf Tip−Ligule distance (cm)
Days
Max. Phytomer Growth Rate
Complete
Simplified
Outliers
adj. R2: −0.3
slope: 2.035
k
Phytomer no.
Rate of Phytomer Deposition
Complete
Outliers
adj. R2: 0.95
slope: 0.417
Phytomer no.
xo (days)
Max. Leaf Distances
Complete
Simplified
Gaussian
Outliers
Linear adj. R2: 0.061
slope: 0.194
L
Phytomer no.

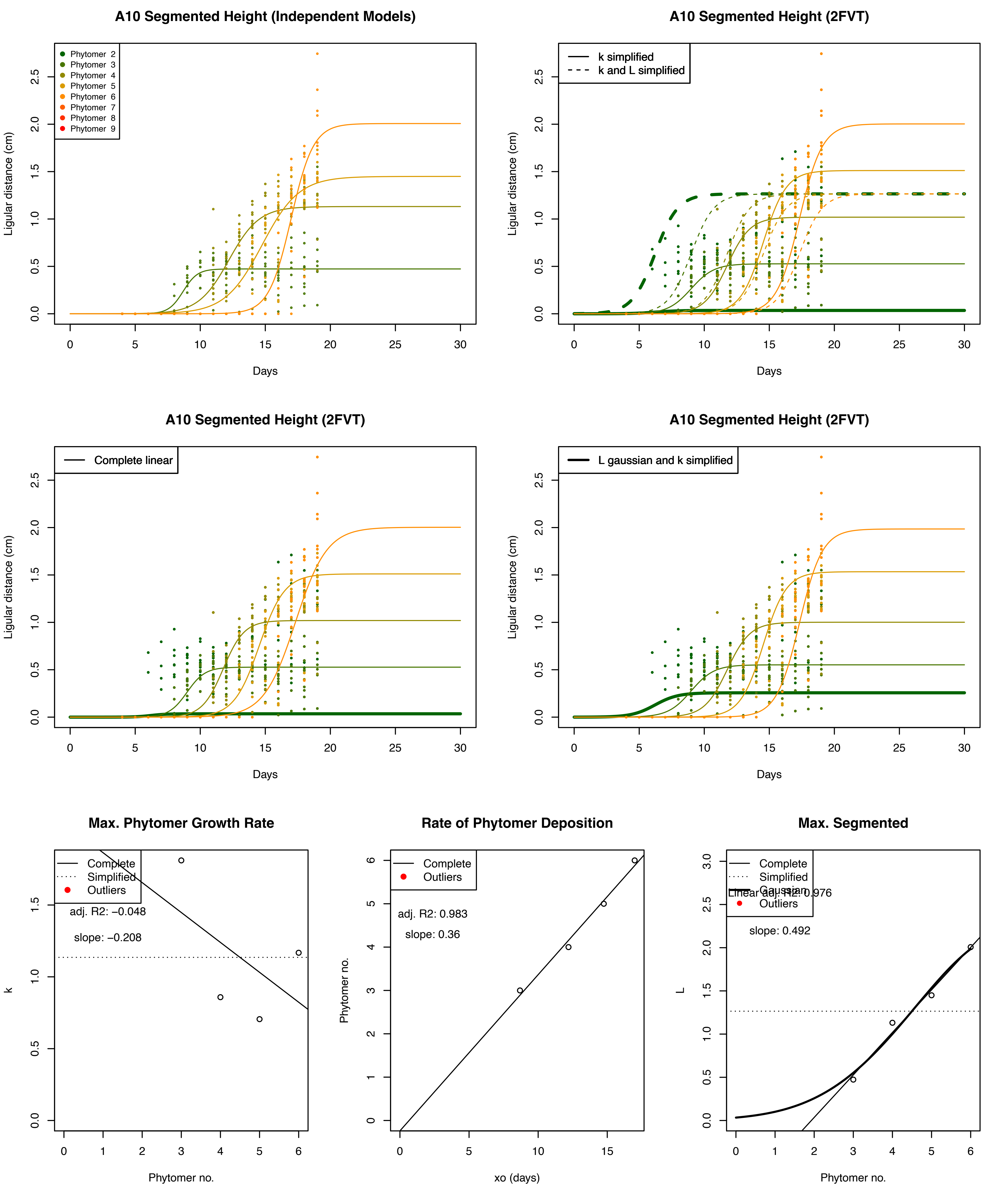

A10 Segmented Height (Independent Models)
Phytomer 2
Phytomer 3
Phytomer 4
Phytomer 5
Phytomer 6
Phytomer 7
Phytomer 8
Phytomer 9
Ligular distance (cm)
Days
A10 Segmented Height (2FVT)
k simplified
k and L simplified
Ligular distance (cm)
Days
A10 Segmented Height (2FVT)
Complete linear
Ligular distance (cm)
Days
A10 Segmented Height (2FVT)
L gaussian and k simplified
Ligular distance (cm)
Days
Max. Phytomer Growth Rate
Complete
Simplified
Outliers
adj. R2: −0.048
slope: −0.208
k
Phytomer no.
Rate of Phytomer Deposition
Complete
Outliers
adj. R2: 0.983
slope: 0.36
Phytomer no.
xo (days)
Max. Segmented
Complete
Simplified
Gaussian
Linear adj. R2: 0.976
Outliers
slope: 0.492
L
Phytomer no.

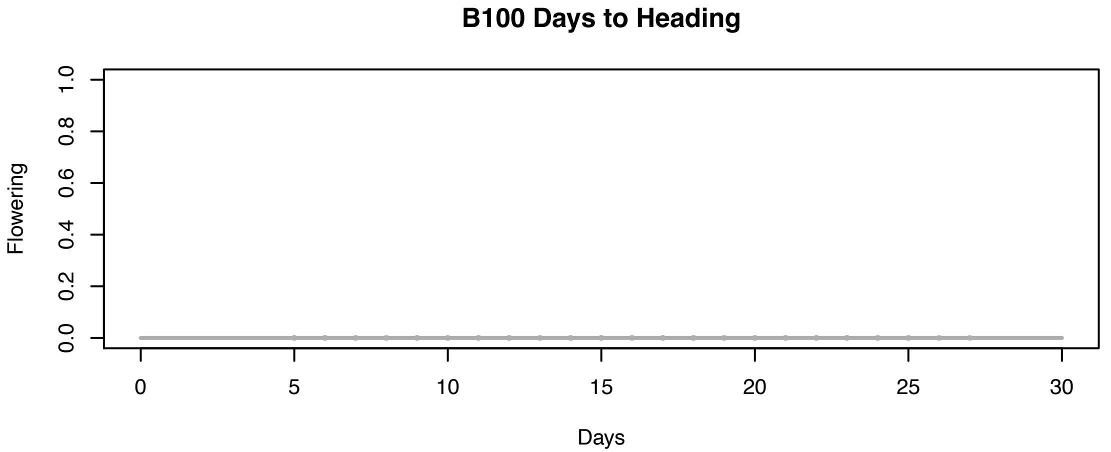
B100 Days to Heading
Flowering
0.0
0.2
0.4
0.6
0.8
1.0
0
5
10
15
20
25
30
Days

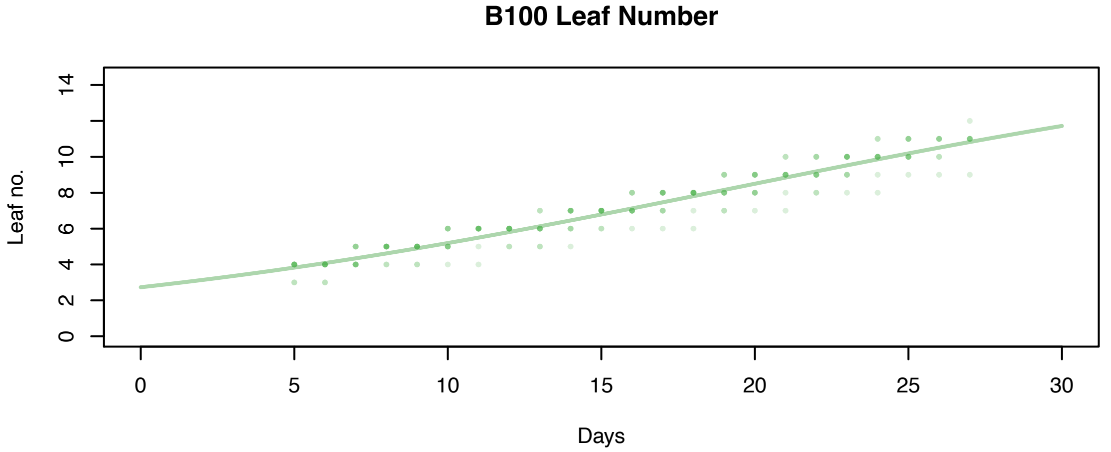
B100 Leaf Number
Leaf no.
0
2
4
6
8
10
14
0
5
10
15
20
25
30
Days

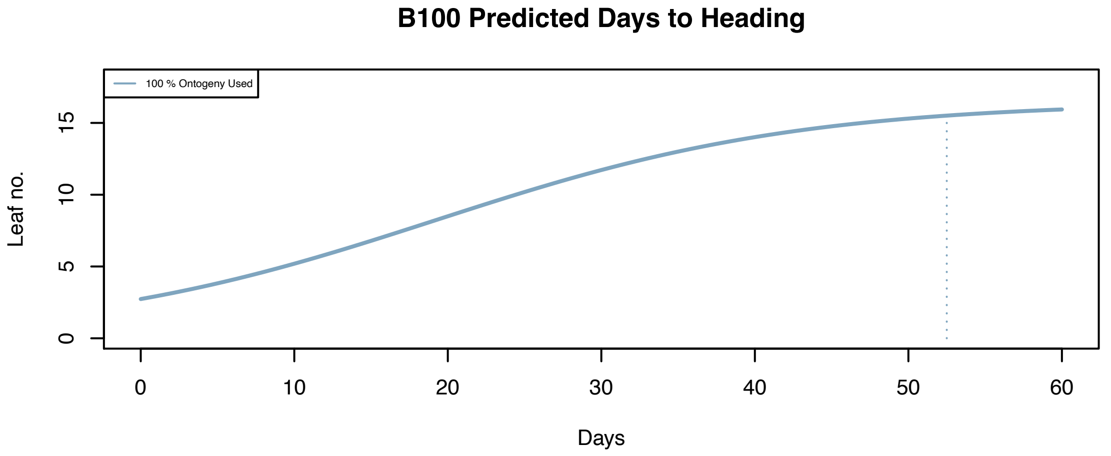
B100 Predicted Days to Heading
100 % Ontogeny Used
Leaf no.
0
5
10
15
0
10
20
30
40
50
60
Days

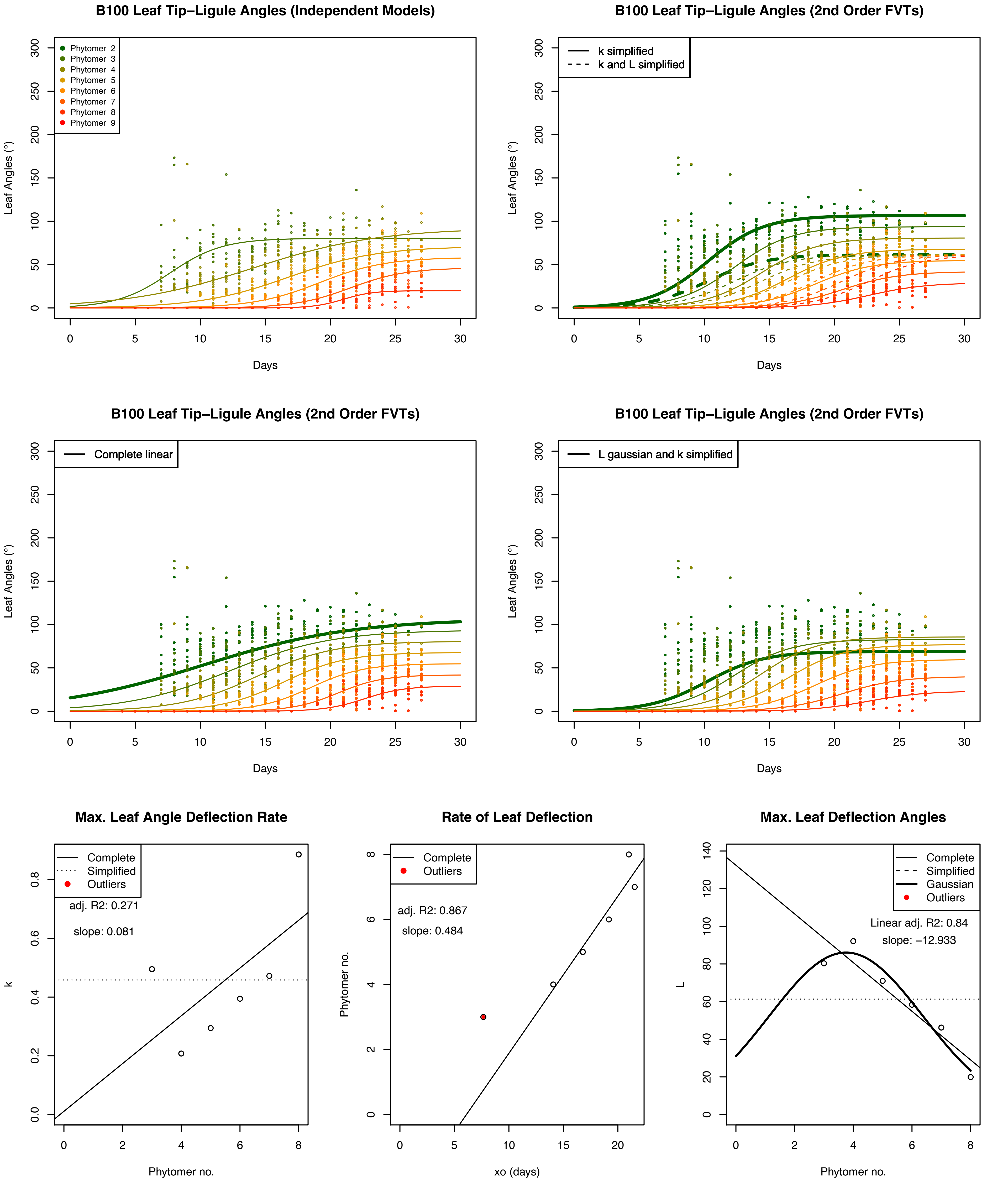

B100 Leaf Tip−Ligule Angles (Independent Models)
Phytomer 2
Phytomer 3
Phytomer 4
Phytomer 5
Phytomer 6
Phytomer 7
Phytomer 8
Phytomer 9
Leaf Angles (°)
Days
B100 Leaf Tip−Ligule Angles (2nd Order FVTs)
k simplified
k and L simplified
Leaf Angles (°)
Days
B100 Leaf Tip−Ligule Angles (2nd Order FVTs)
Complete linear
Leaf Angles (°)
Days
B100 Leaf Tip−Ligule Angles (2nd Order FVTs)
L gaussian and k simplified
Leaf Angles (°)
Days
Max. Leaf Angle Deflection Rate
Complete
Simplified
Outliers
adj. R2: 0.271
slope: 0.081
k
Phytomer no.
Rate of Leaf Deflection
Complete
Outliers
adj. R2: 0.867
slope: 0.484
Phytomer no.
xo (days)
Max. Leaf Deflection Angles
Complete
Simplified
Gaussian
Outliers
Linear adj. R2: 0.84
slope: −12.933
L
Phytomer no.

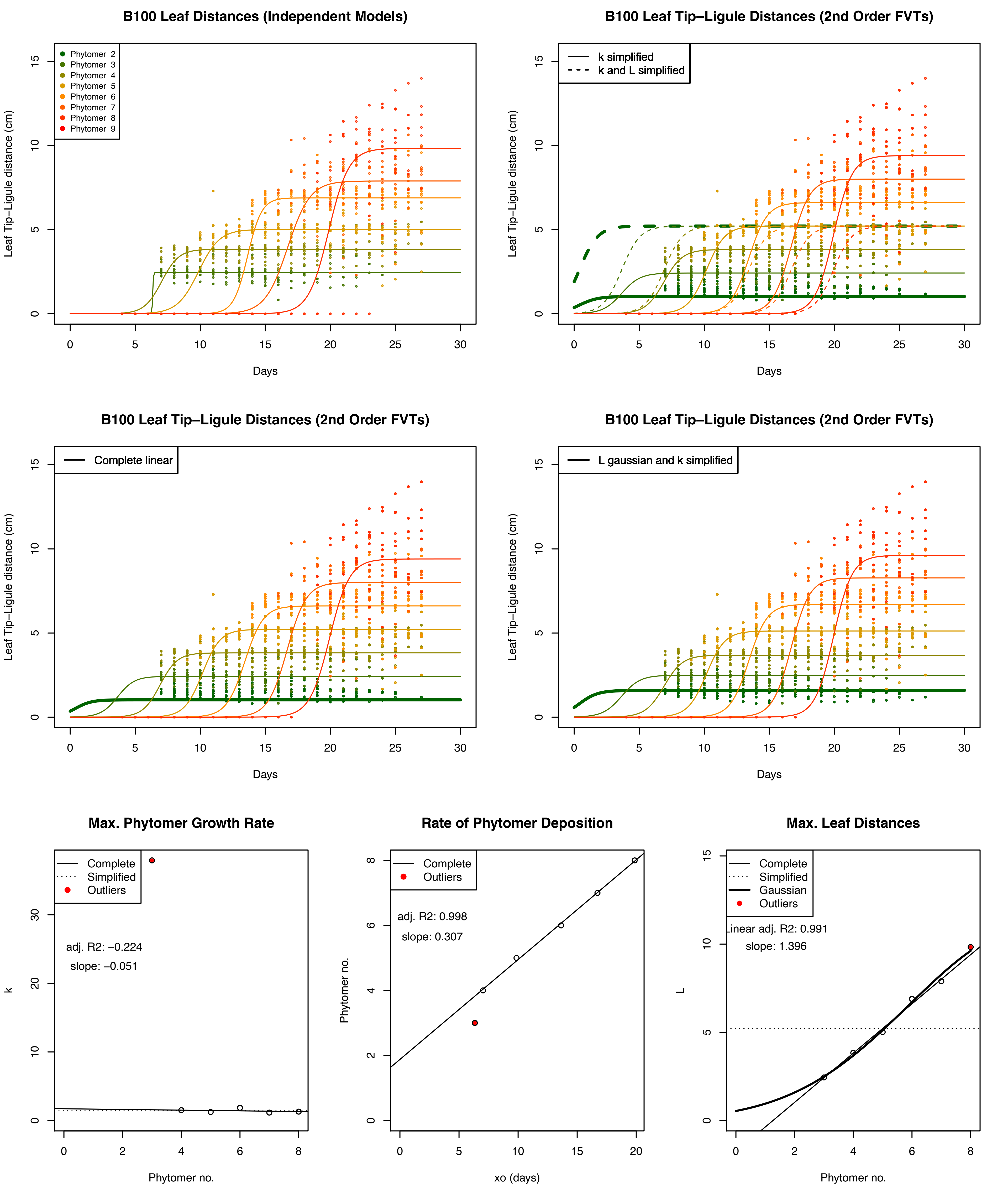
B100 Leaf Distances (Independent Models)
Phytomer 2
Phytomer 3
Phytomer 4
Phytomer 5
Phytomer 6
Phytomer 7
Phytomer 8
Phytomer 9
Leaf Tip−Ligule distance (cm)
Days
B100 Leaf Tip−Ligule Distances (2nd Order FVTs)
k simplified
k and L simplified
Leaf Tip−Ligule distance (cm)
Days
B100 Leaf Tip−Ligule Distances (2nd Order FVTs)
Complete linear
Leaf Tip−Ligule distance (cm)
Days
B100 Leaf Tip−Ligule Distances (2nd Order FVTs)
L gaussian and k simplified
Leaf Tip−Ligule distance (cm)
Days
Max. Phytomer Growth Rate
Complete
Simplified
Outliers
adj. R2: −0.224
slope: −0.051
k
Phytomer no.
Rate of Phytomer Deposition
Complete
Outliers
adj. R2: 0.998
slope: 0.307
Phytomer no.
xo (days)
Max. Leaf Distances
Complete
Simplified
Gaussian
Outliers
Linear adj. R2: 0.991
slope: 1.396
L
Phytomer no.

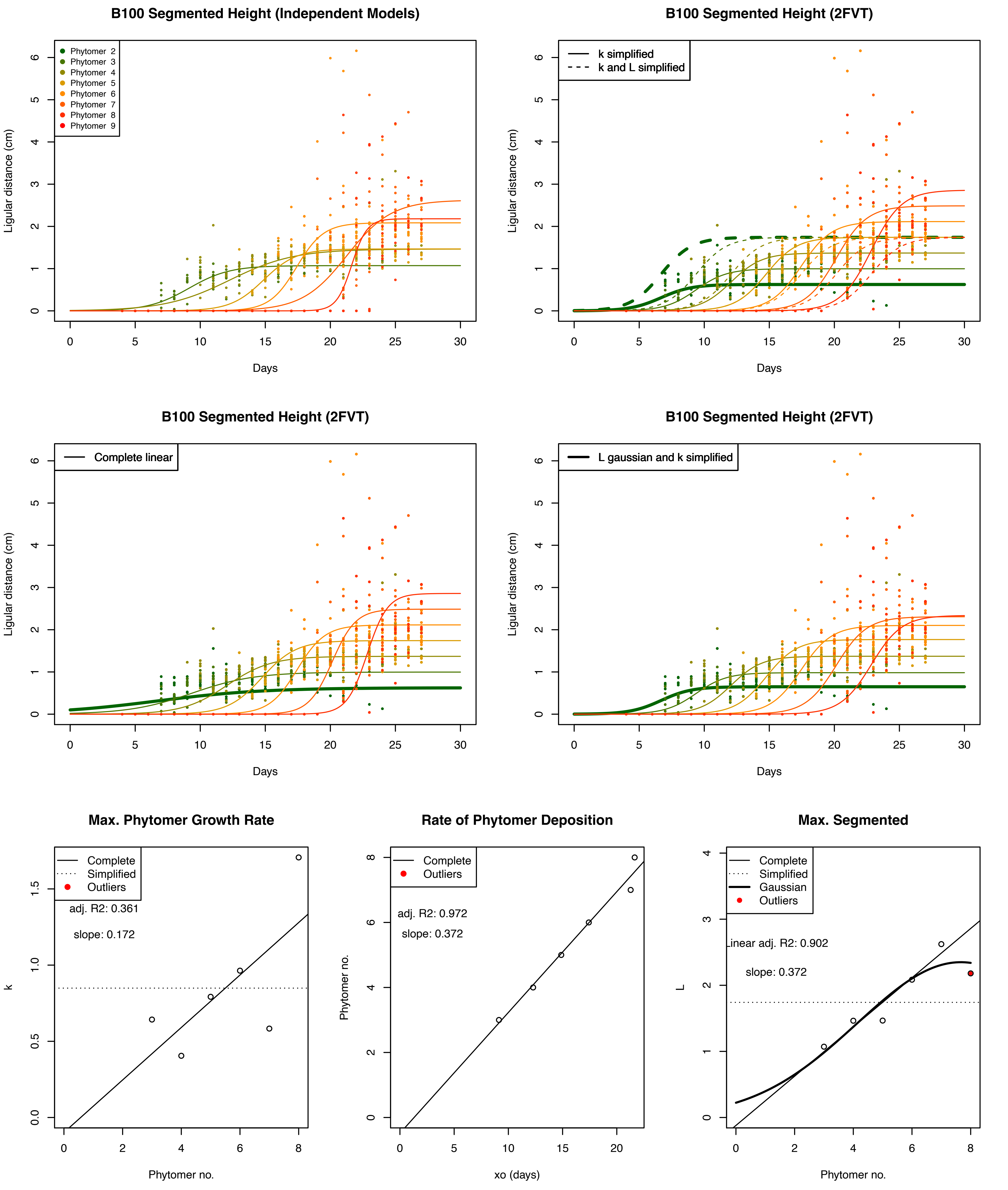
B100 Segmented Height (Independent Models)
Phytomer 2
Phytomer 3
Phytomer 4
Phytomer 5
Phytomer 6
Phytomer 7
Phytomer 8
Phytomer 9
Ligular distance (cm)
Days
B100 Segmented Height (2FVT)
k simplified
k and L simplified
Ligular distance (cm)
Days
B100 Segmented Height (2FVT)
Complete linear
Ligular distance (cm)
Days
B100 Segmented Height (2FVT)
L gaussian and k simplified
Ligular distance (cm)
Days
Max. Phytomer Growth Rate
Complete
Simplified
Outliers
adj. R2: 0.361
slope: 0.172
k
Phytomer no.
Rate of Phytomer Deposition
Complete
Outliers
adj. R2: 0.972
slope: 0.372
Phytomer no.
xo (days)
Max. Segmented
Complete
Simplified
Gaussian
Outliers
Linear adj. R2: 0.902
slope: 0.372
L
Phytomer no.

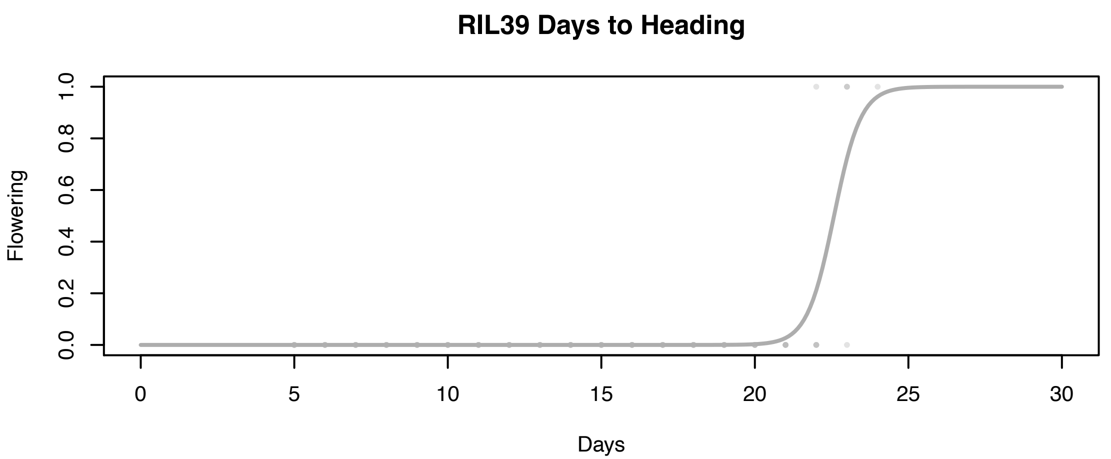
RIL39 Days to Heading
Flowering
0.0
0.2
0.4
0.6
0.8
1.0
0
5
10
15
20
25
30
Days

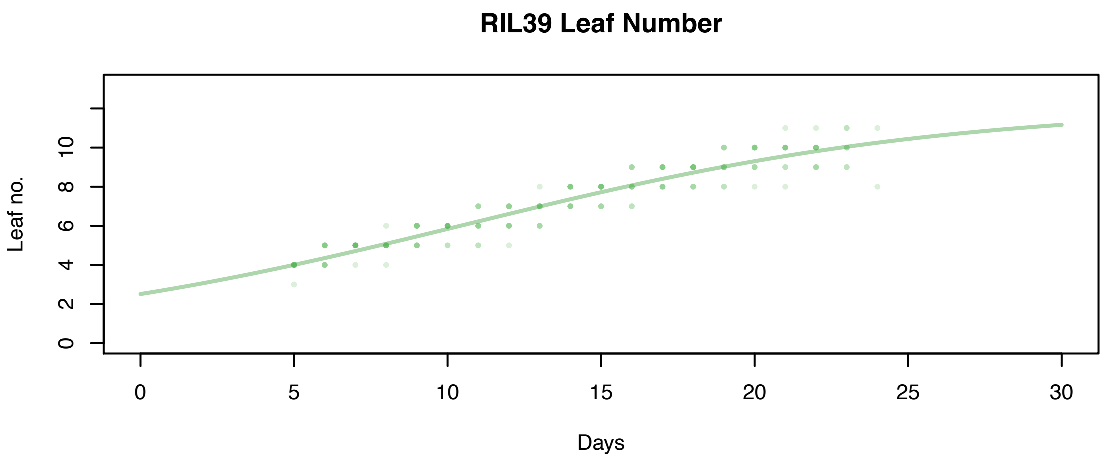
RIL39 Leaf Number
Leaf no.
0
2
4
6
8
10
0
5
10
15
20
25
30
Days

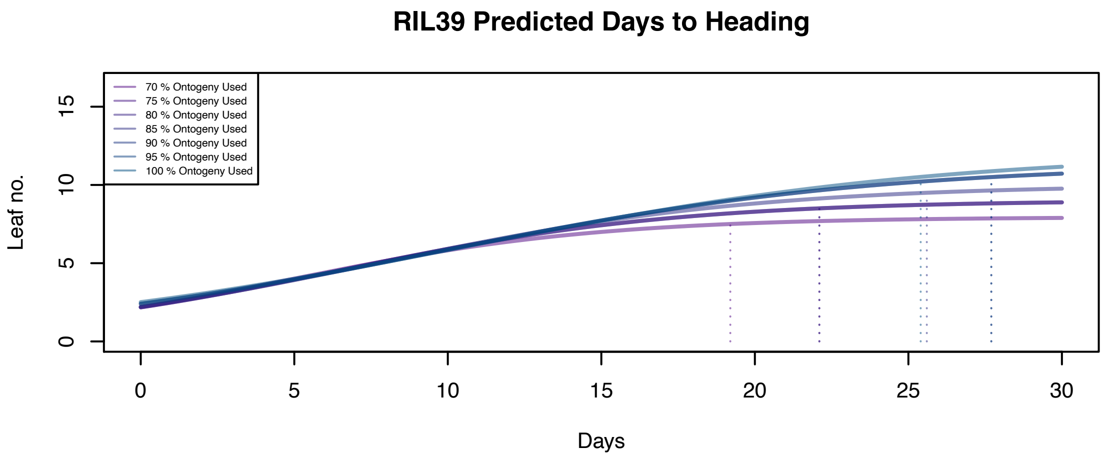
RIL39 Predicted Days to Heading
70 % Ontogeny Used
75 % Ontogeny Used
80 % Ontogeny Used
85 % Ontogeny Used
90 % Ontogeny Used
95 % Ontogeny Used
100 % Ontogeny Used
Leaf no.
0
5
10
15
0
5
10
15
20
25
30
Days

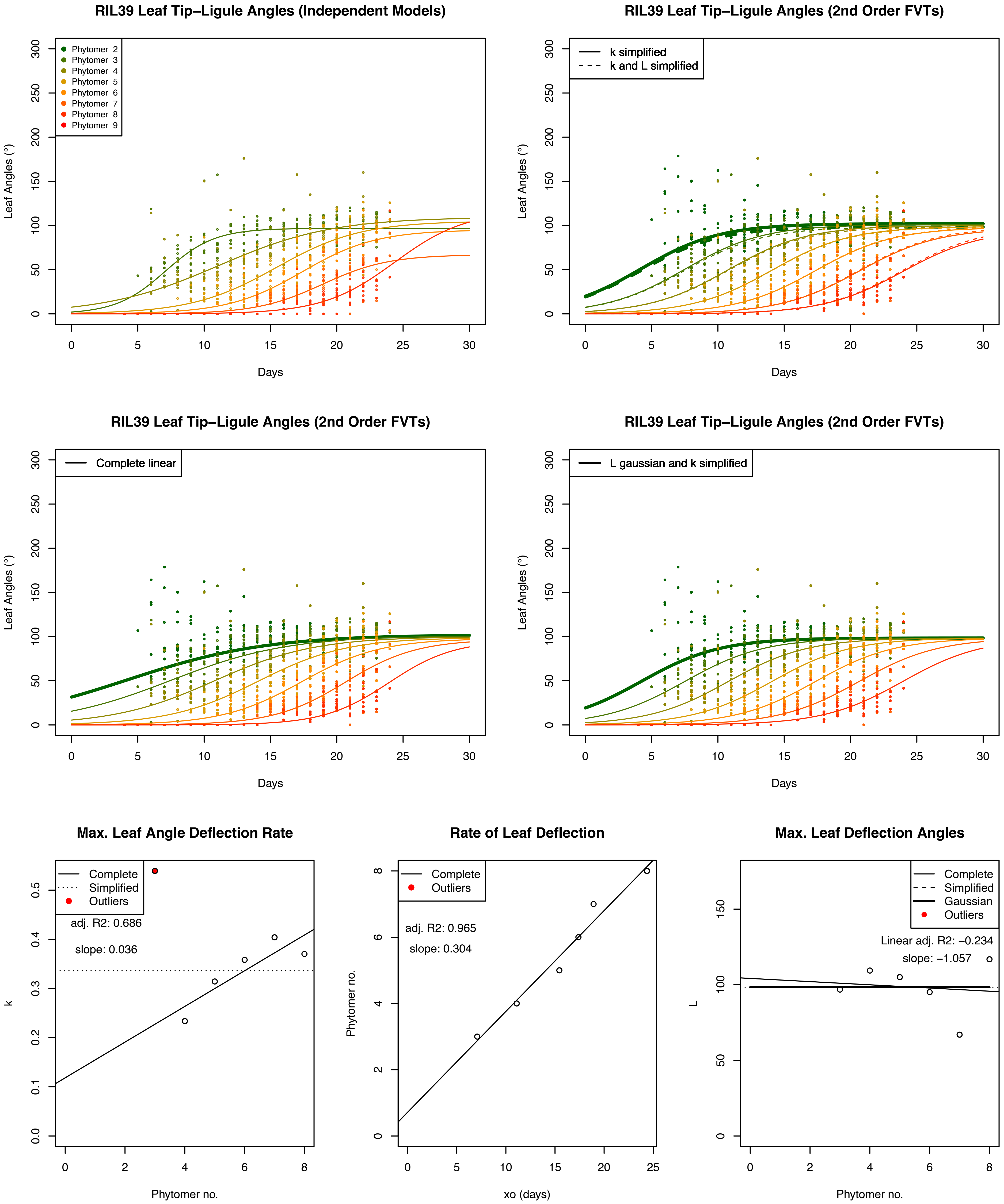
RIL39 Leaf Tip–Ligule Angles (Independent Models)
Phytomer 2
Phytomer 3
Phytomer 4
Phytomer 5
Phytomer 6
Phytomer 7
Phytomer 8
Phytomer 9
Leaf Angles (°)
Days
RIL39 Leaf Tip–Ligule Angles (2nd Order FVTs)
k simplified
k and L simplified
Leaf Angles (°)
Days
RIL39 Leaf Tip–Ligule Angles (2nd Order FVTs)
Complete linear
Leaf Angles (°)
Days
RIL39 Leaf Tip–Ligule Angles (2nd Order FVTs)
L gaussian and k simplified
Leaf Angles (°)
Days
Max. Leaf Angle Deflection Rate
Complete
Simplified
Outliers
adj. R2: 0.686
slope: 0.036
k
Phytomer no.
Rate of Leaf Deflection
Complete
Outliers
adj. R2: 0.965
slope: 0.304
Phytomer no.
xo (days)
Max. Leaf Deflection Angles
Complete
Simplified
Gaussian
Outliers
Linear adj. R2: −0.234
slope: −1.057
L
Phytomer no.

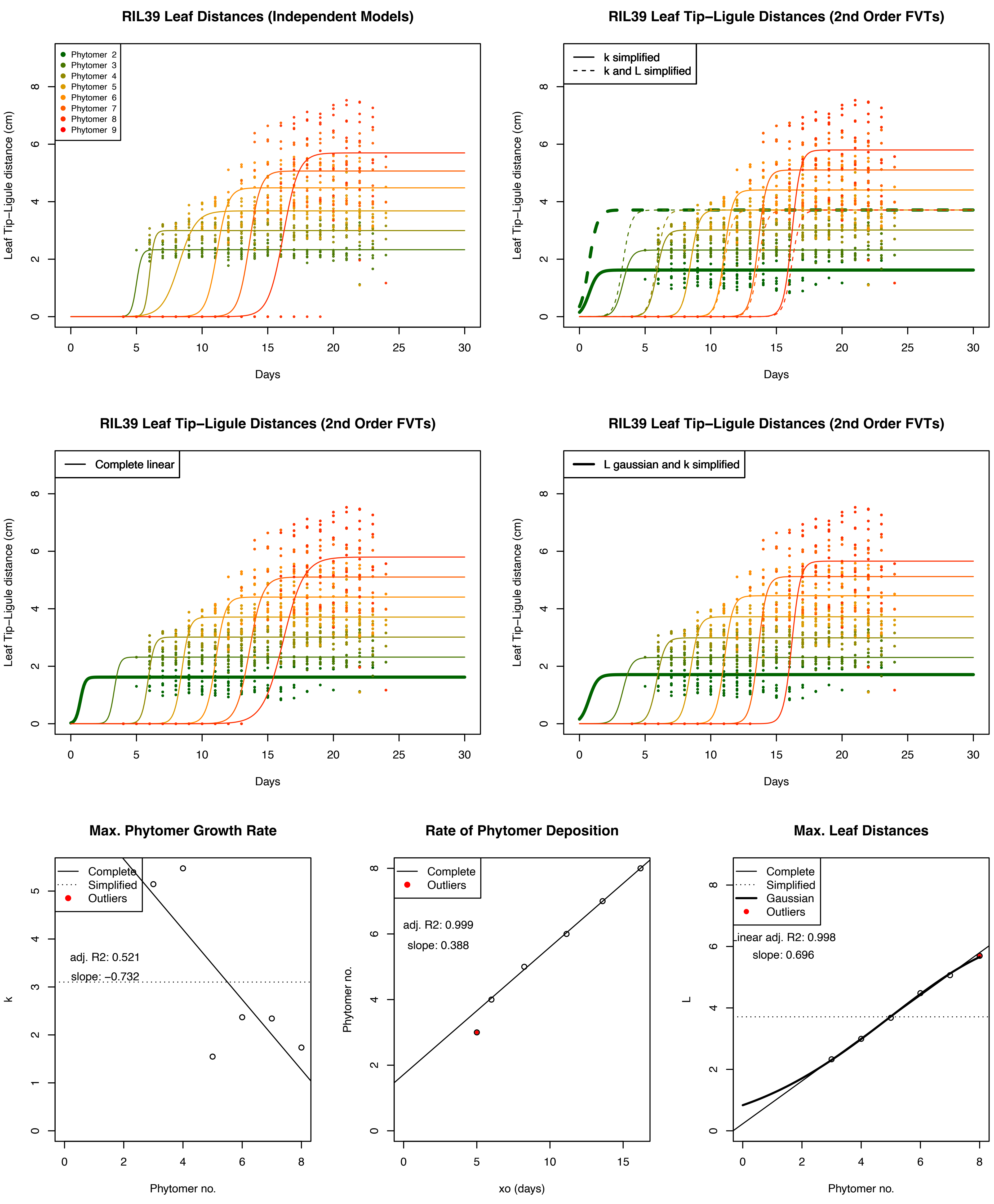
RIL39 Leaf Distances (Independent Models)
Phytomer 2
Phytomer 3
Phytomer 4
Phytomer 5
Phytomer 6
Phytomer 7
Phytomer 8
Phytomer 9
Leaf Tip−Ligule distance (cm)
Days
RIL39 Leaf Tip−Ligule Distances (2nd Order FVTs)
k simplified
k and L simplified
Leaf Tip−Ligule distance (cm)
Days
RIL39 Leaf Tip−Ligule Distances (2nd Order FVTs)
Complete linear
Leaf Tip−Ligule distance (cm)
Days
RIL39 Leaf Tip−Ligule Distances (2nd Order FVTs)
L gaussian and k simplified
Leaf Tip−Ligule distance (cm)
Days
Max. Phytomer Growth Rate
Complete
Simplified
Outliers
adj. R2: 0.521
slope: −0.732
k
Phytomer no.
Rate of Phytomer Deposition
Complete
Outliers
adj. R2: 0.999
slope: 0.388
Phytomer no.
xo (days)
Max. Leaf Distances
Complete
Simplified
Gaussian
Outliers
Linear adj. R2: 0.998
slope: 0.696
L
Phytomer no.

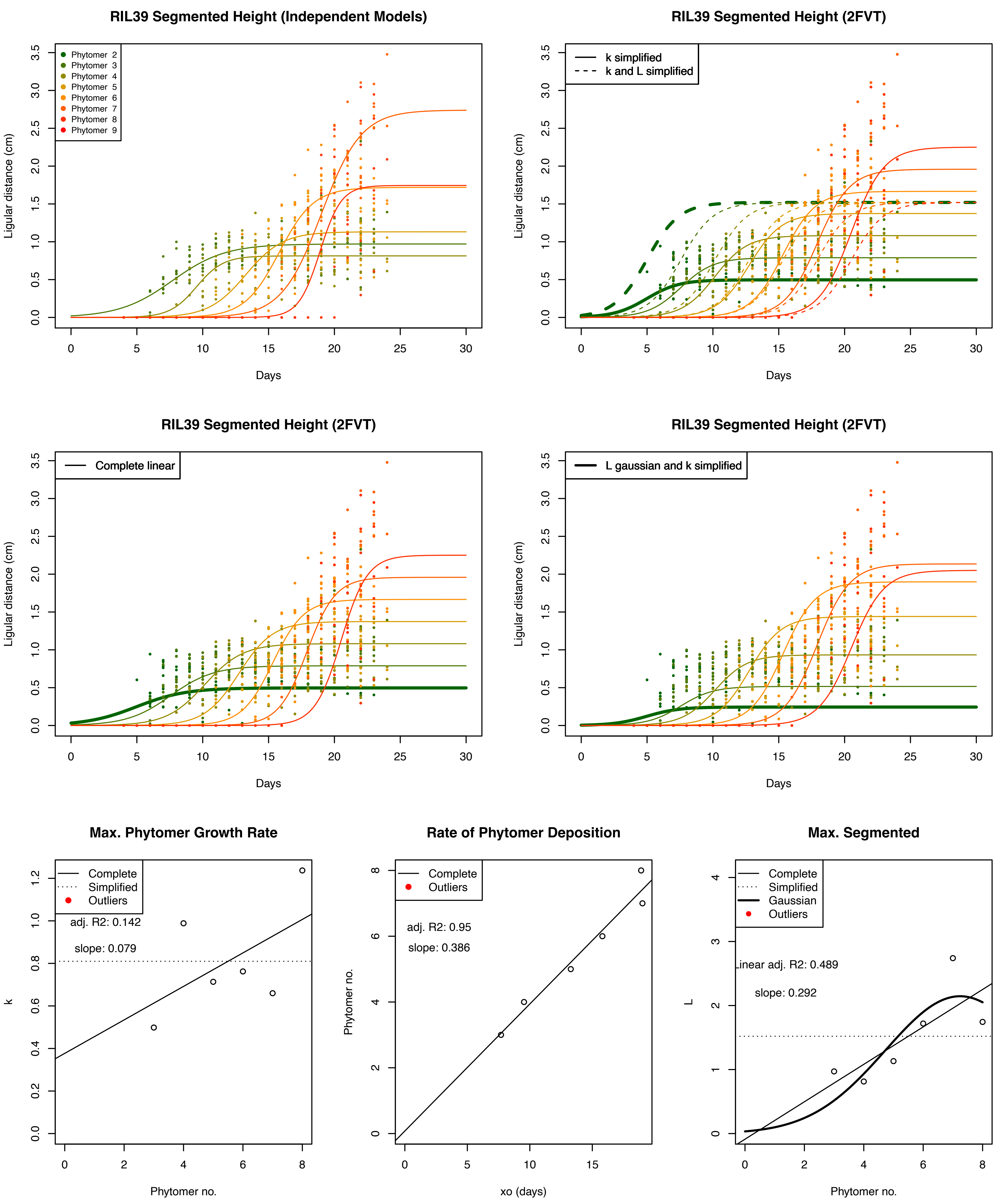
RIL39 Segmented Height (Independent Models)
Phytomer 2
Phytomer 3
Phytomer 4
Phytomer 5
Phytomer 6
Phytomer 7
Phytomer 8
Phytomer 9
Ligular distance (cm)
Days
RIL39 Segmented Height (2FVT)
k simplified
k and L simplified
Ligular distance (cm)
Days
RIL39 Segmented Height (2FVT)
Complete linear
Ligular distance (cm)
Days
RIL39 Segmented Height (2FVT)
L gaussian and k simplified
Ligular distance (cm)
Days
Max. Phytomer Growth Rate
Complete
Simplified
Outliers
adj. R2: 0.142
slope: 0.079
k
Phytomer no.
Rate of Phytomer Deposition
Complete
Outliers
adj. R2: 0.95
slope: 0.386
Phytomer no.
xo (days)
Max. Segmented
Complete
Simplified
Gaussian
Outliers
Linear adj. R2: 0.489
slope: 0.292
L
Phytomer no.

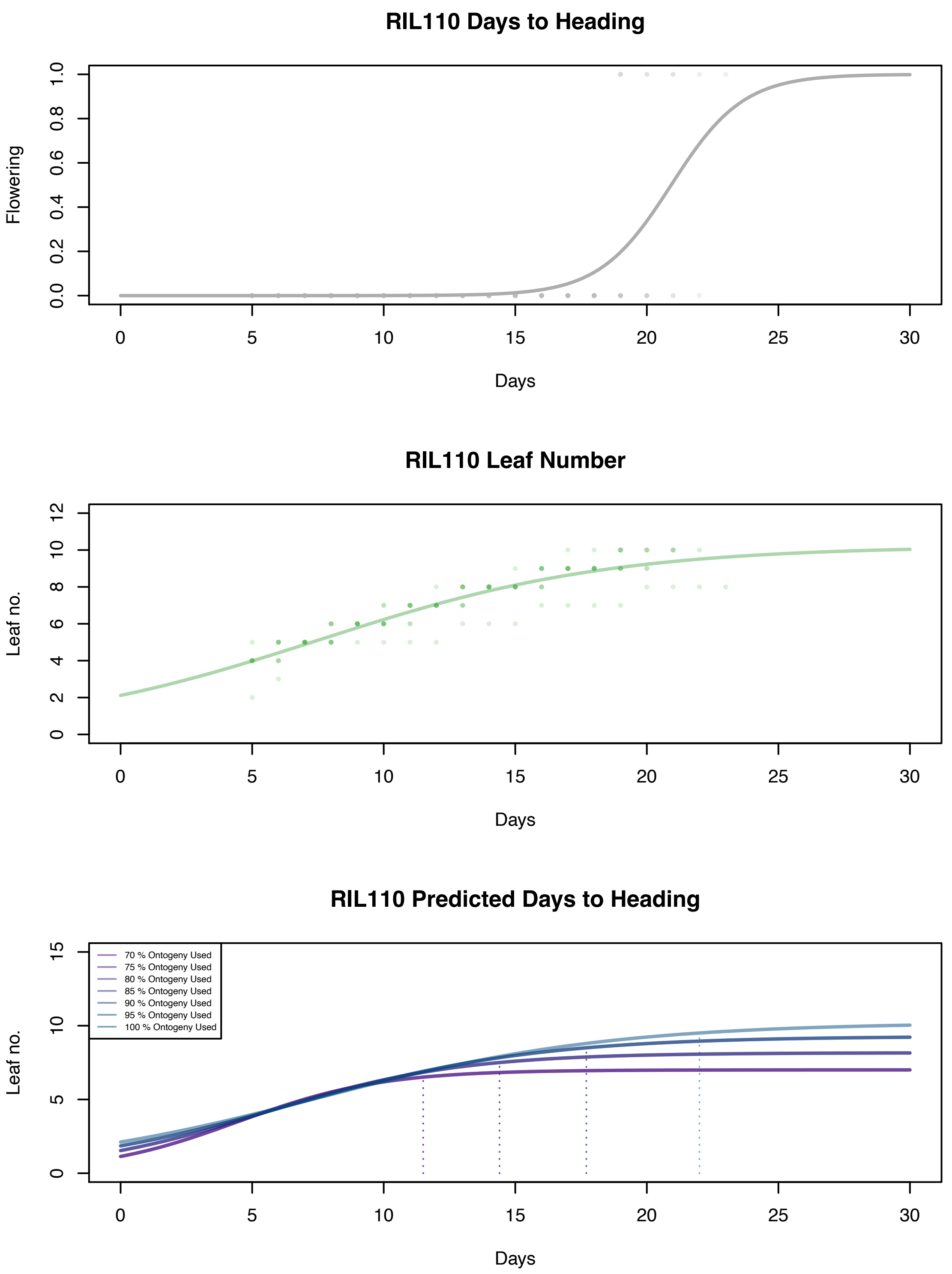
RIL110 Days to Heading
Flowering
0.0
0.2
0.4
0.6
0.8
1.0
0
5
10
15
20
25
30
Days
RIL110 Leaf Number
Leaf no.
0
2
4
6
8
10
12
Days
RIL110 Predicted Days to Heading
Leaf no.
0
5
10
15
70 % Ontogeny Used
75 % Ontogeny Used
80 % Ontogeny Used
85 % Ontogeny Used
90 % Ontogeny Used
95 % Ontogeny Used
100 % Ontogeny Used
Days

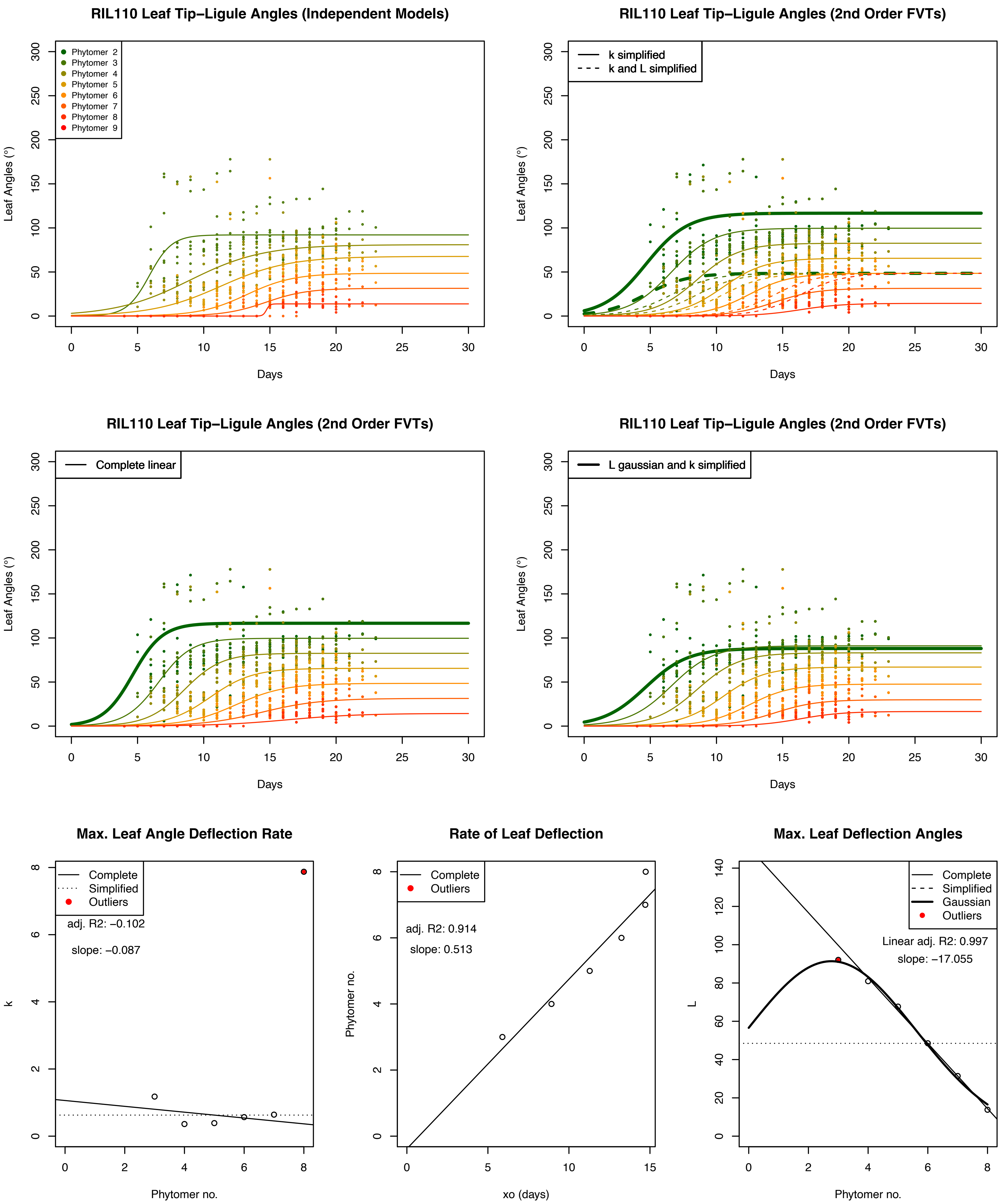

RIL110 Leaf Tip–Ligule Angles (Independent Models)
Phytomer 2
Phytomer 3
Phytomer 4
Phytomer 5
Phytomer 6
Phytomer 7
Phytomer 8
Phytomer 9
Leaf Angles (°)
Days
RIL110 Leaf Tip–Ligule Angles (2nd Order FVTs)
k simplified
k and L simplified
Leaf Angles (°)
Days
RIL110 Leaf Tip–Ligule Angles (2nd Order FVTs)
Complete linear
Leaf Angles (°)
Days
RIL110 Leaf Tip–Ligule Angles (2nd Order FVTs)
L gaussian and k simplified
Leaf Angles (°)
Days
Max. Leaf Angle Deflection Rate
Complete
Simplified
Outliers
adj. R2: −0.102
slope: −0.087
k
Phytomer no.
Rate of Leaf Deflection
Complete
Outliers
adj. R2: 0.914
slope: 0.513
Phytomer no.
xo (days)
Max. Leaf Deflection Angles
Complete
Simplified
Gaussian
Outliers
Linear adj. R2: 0.997
slope: −17.055
L
Phytomer no.

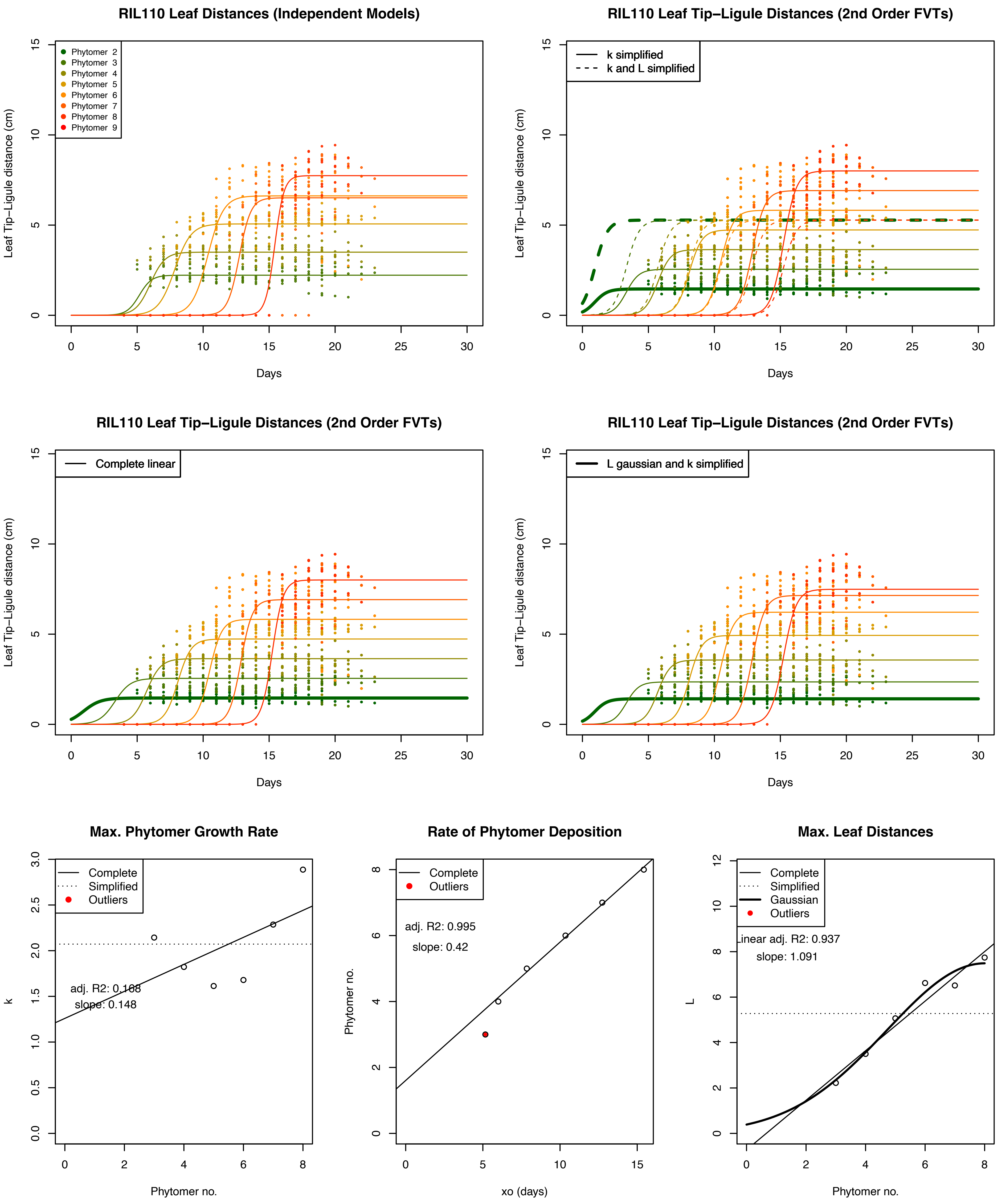
RIL110 Leaf Distances (Independent Models)
Leaf Tip−Ligule distance (cm)
Phytomer 2
Phytomer 3
Phytomer 4
Phytomer 5
Phytomer 6
Phytomer 7
Phytomer 8
Phytomer 9
Days
RIL110 Leaf Tip−Ligule Distances (2nd Order FVTs)
k simplified
k and L simplified
Leaf Tip−Ligule distance (cm)
Days
RIL110 Leaf Tip−Ligule Distances (2nd Order FVTs)
Complete linear
Leaf Tip−Ligule distance (cm)
Days
RIL110 Leaf Tip−Ligule Distances (2nd Order FVTs)
L gaussian and k simplified
Leaf Tip−Ligule distance (cm)
Days
Max. Phytomer Growth Rate
Complete
Simplified
Outliers
adj. R2: 0.168
slope: 0.148
k
Phytomer no.
Rate of Phytomer Deposition
Complete
Outliers
adj. R2: 0.995
slope: 0.42
Phytomer no.
xo (days)
Max. Leaf Distances
Complete
Simplified
Gaussian
Outliers
Linear adj. R2: 0.937
slope: 1.091
L
Phytomer no.

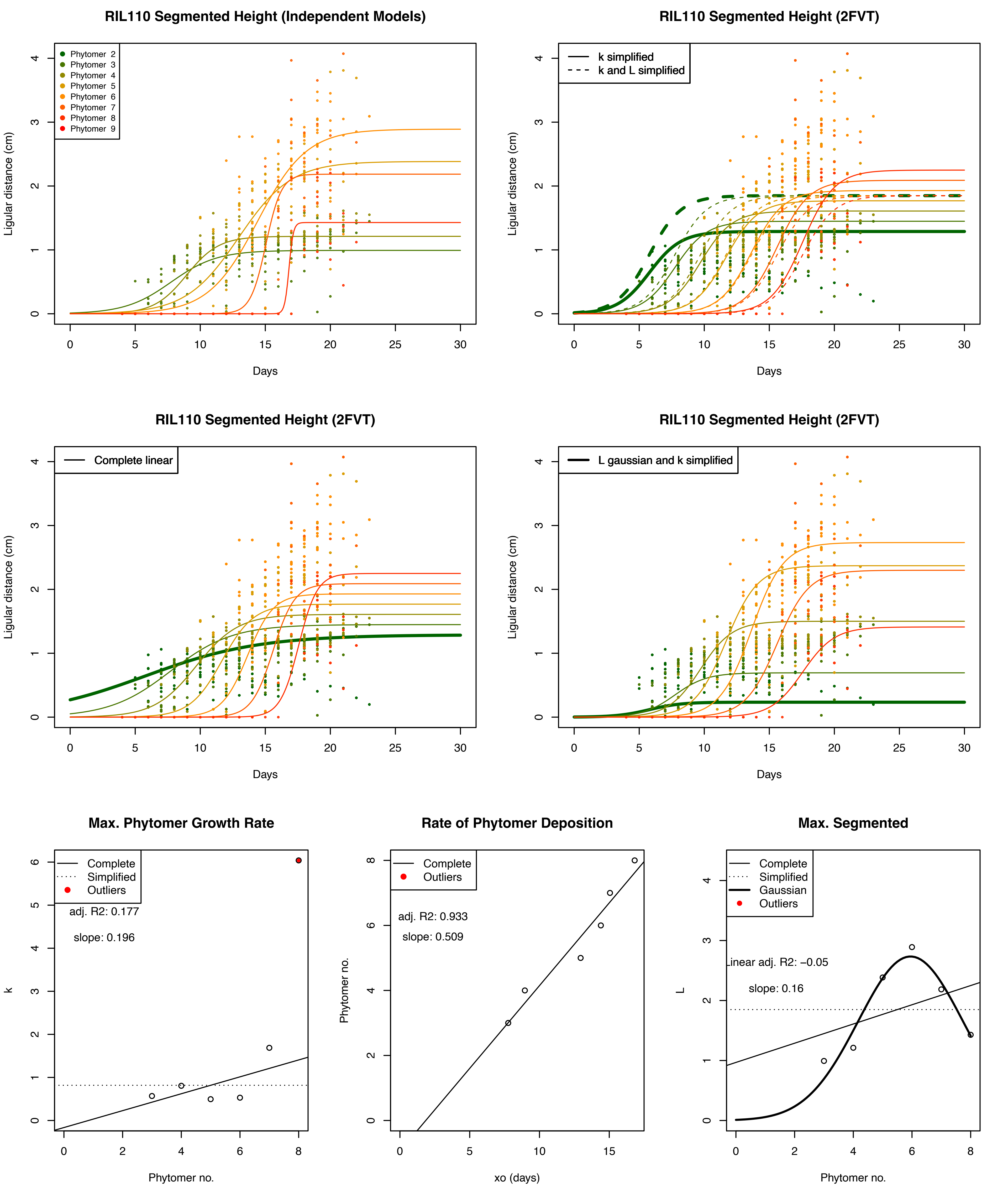

RIL110 Segmented Height (Independent Models)
Phytomer 2
Phytomer 3
Phytomer 4
Phytomer 5
Phytomer 6
Phytomer 7
Phytomer 8
Phytomer 9
Ligular distance (cm)
Days
RIL110 Segmented Height (2FVT)
k simplified
k and L simplified
Ligular distance (cm)
Days
RIL110 Segmented Height (2FVT)
Complete linear
Ligular distance (cm)
Days
RIL110 Segmented Height (2FVT)
L gaussian and k simplified
Ligular distance (cm)
Days
Max. Phytomer Growth Rate
Complete
Simplified
Outliers
adj. R2: 0.177
slope: 0.196
k
Phytomer no.
Rate of Phytomer Deposition
Complete
Outliers
adj. R2: 0.933
slope: 0.509
Phytomer no.
xo (days)
Max. Segmented
Complete
Simplified
Gaussian
Outliers
Linear adj. R2: −0.05
slope: 0.16
L
Phytomer no.

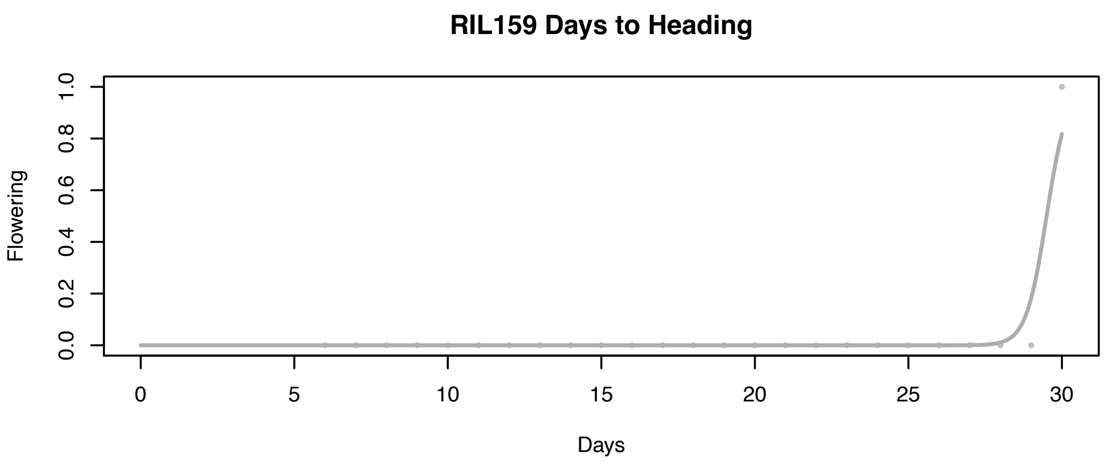

RIL159 Days to Heading
Flowering
0.0
0.2
0.4
0.6
0.8
1.0
0
5
10
15
20
25
30
Days


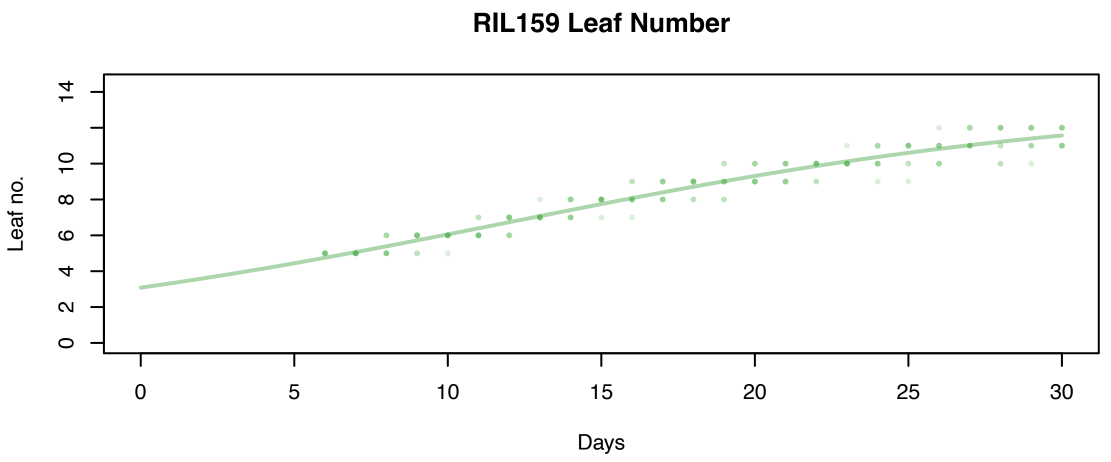

RIL159 Leaf Number
Leaf no.
0
2
4
6
8
10
14
0
5
10
15
20
25
30
Days


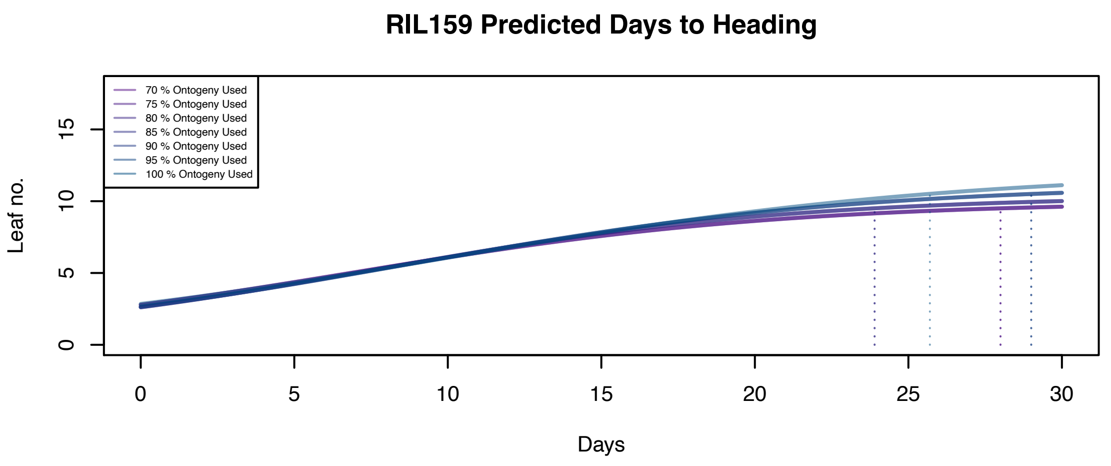

RIL159 Predicted Days to Heading
70 % Ontogeny Used
75 % Ontogeny Used
80 % Ontogeny Used
85 % Ontogeny Used
90 % Ontogeny Used
95 % Ontogeny Used
100 % Ontogeny Used
Leaf no.
0
5
10
15
0
5
10
15
20
25
30
Days

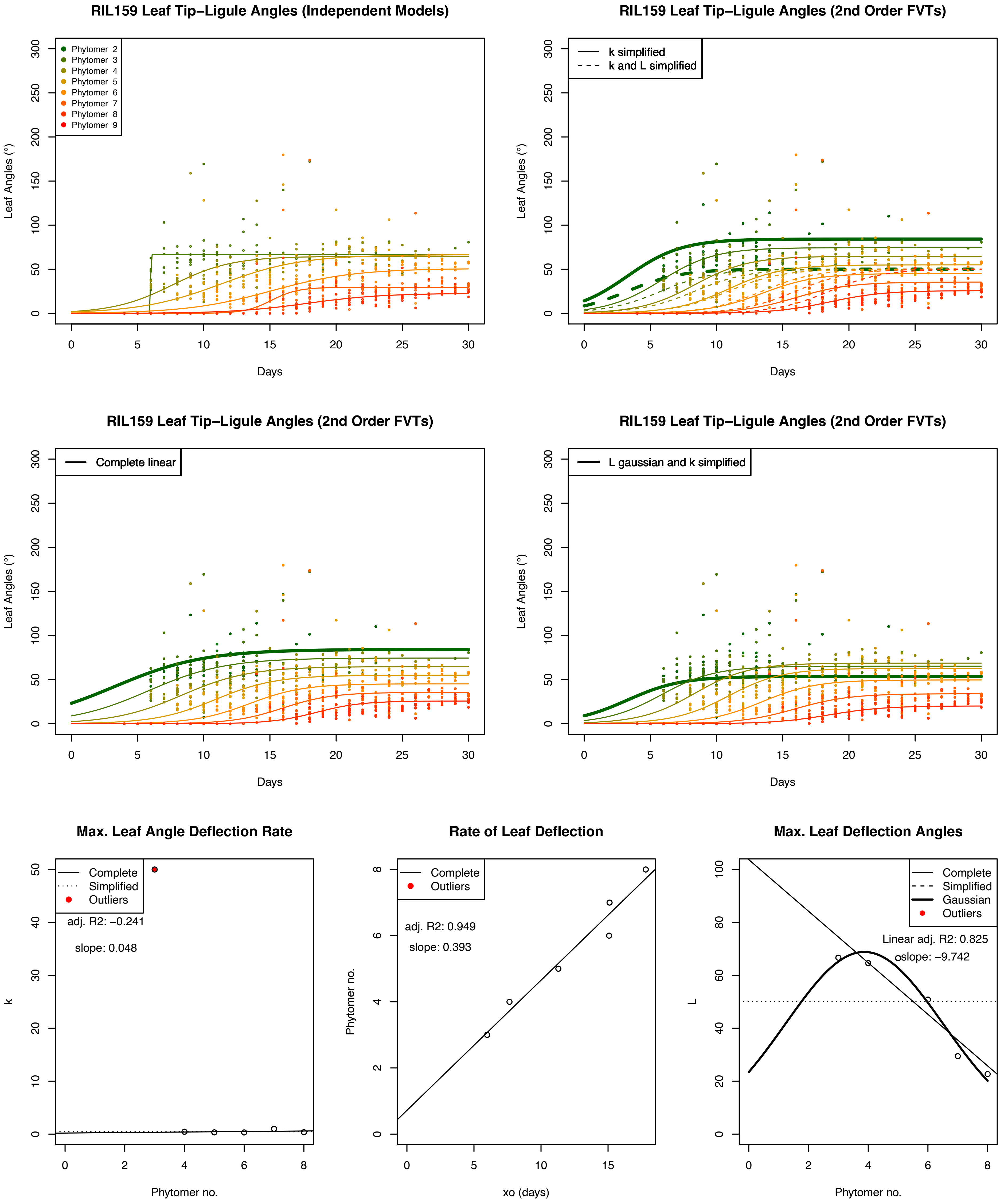
RIL159 Leaf Tip–Ligule Angles (Independent Models)
Phytomer 2
Phytomer 3
Phytomer 4
Phytomer 5
Phytomer 6
Phytomer 7
Phytomer 8
Phytomer 9
Leaf Angles (°)
Days
RIL159 Leaf Tip–Ligule Angles (2nd Order FVTs)
k simplified
k and L simplified
Leaf Angles (°)
Days
RIL159 Leaf Tip–Ligule Angles (2nd Order FVTs)
Complete linear
Leaf Angles (°)
Days
RIL159 Leaf Tip–Ligule Angles (2nd Order FVTs)
L gaussian and k simplified
Leaf Angles (°)
Days
Max. Leaf Angle Deflection Rate
Complete
Simplified
Outliers
adj. R2: –0.241
slope: 0.048
k
Phytomer no.
Rate of Leaf Deflection
Complete
Outliers
adj. R2: 0.949
slope: 0.393
Phytomer no.
xo (days)
Max. Leaf Deflection Angles
Complete
Simplified
Gaussian
Outliers
Linear adj. R2: 0.825
slope: –9.742
L
Phytomer no.

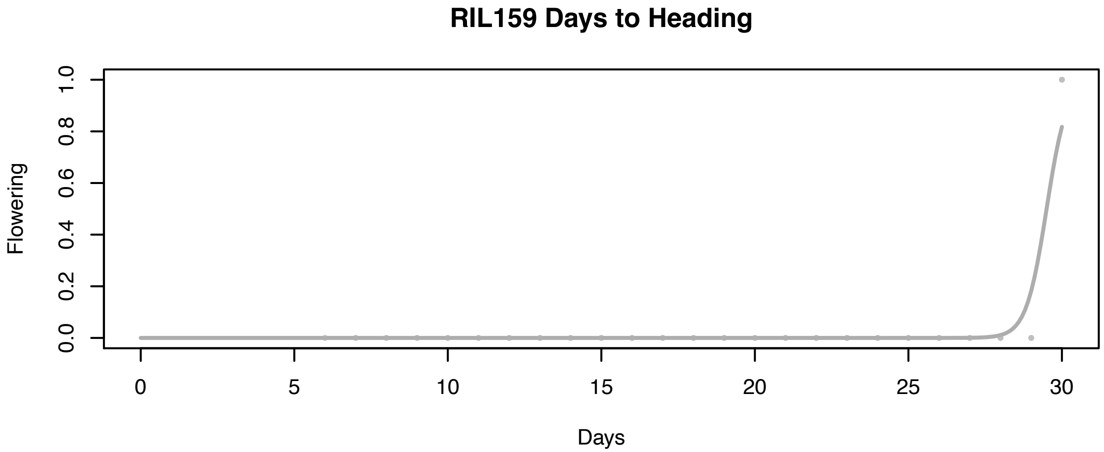
RIL159 Days to Heading
Flowering
Days
0.0
0.2
0.4
0.6
0.8
1.0
0
5
10
15
20
25
30

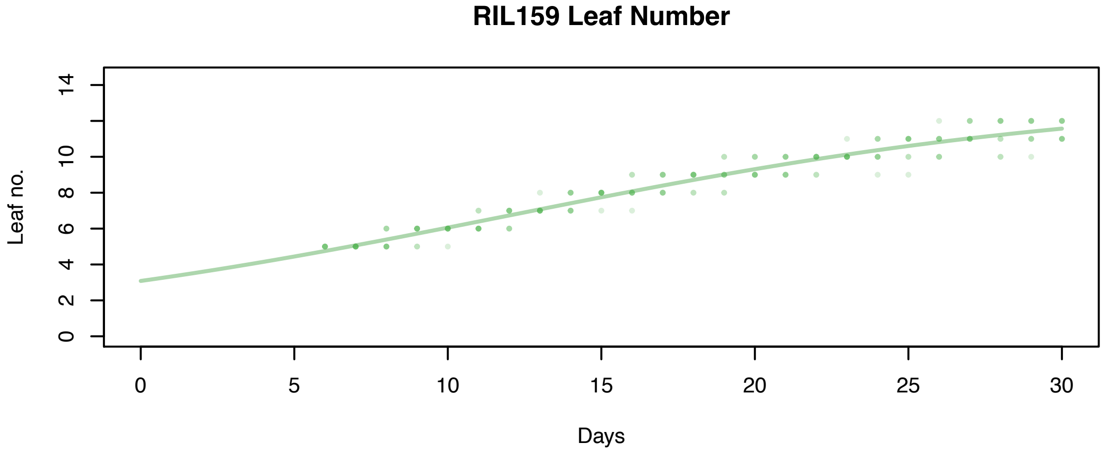
RIL159 Leaf Number
Leaf no.
Days
0
2
4
6
8
10
14
0
5
10
15
20
25
30

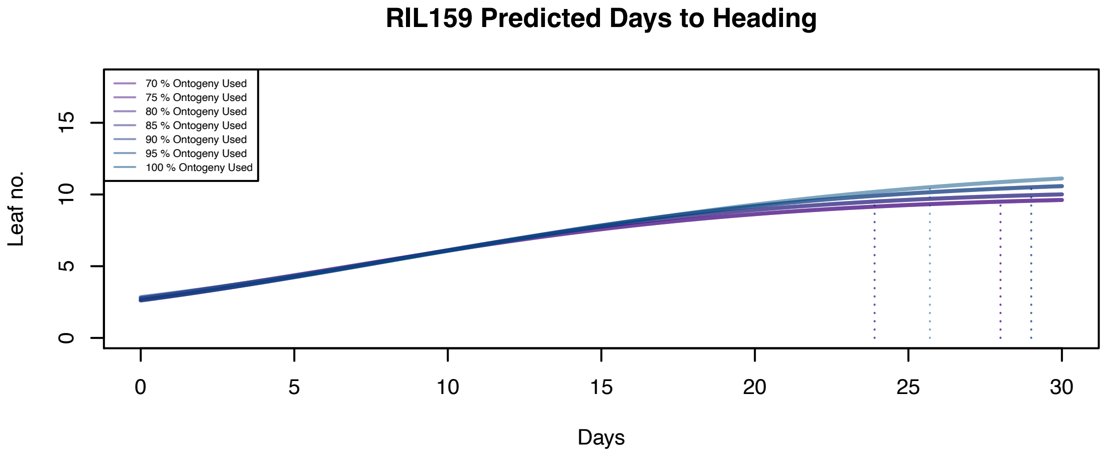
RIL159 Predicted Days to Heading
70 % Ontogeny Used
75 % Ontogeny Used
80 % Ontogeny Used
85 % Ontogeny Used
90 % Ontogeny Used
95 % Ontogeny Used
100 % Ontogeny Used
Leaf no.
Days
0
5
10
15
0
5
10
15
20
25
30

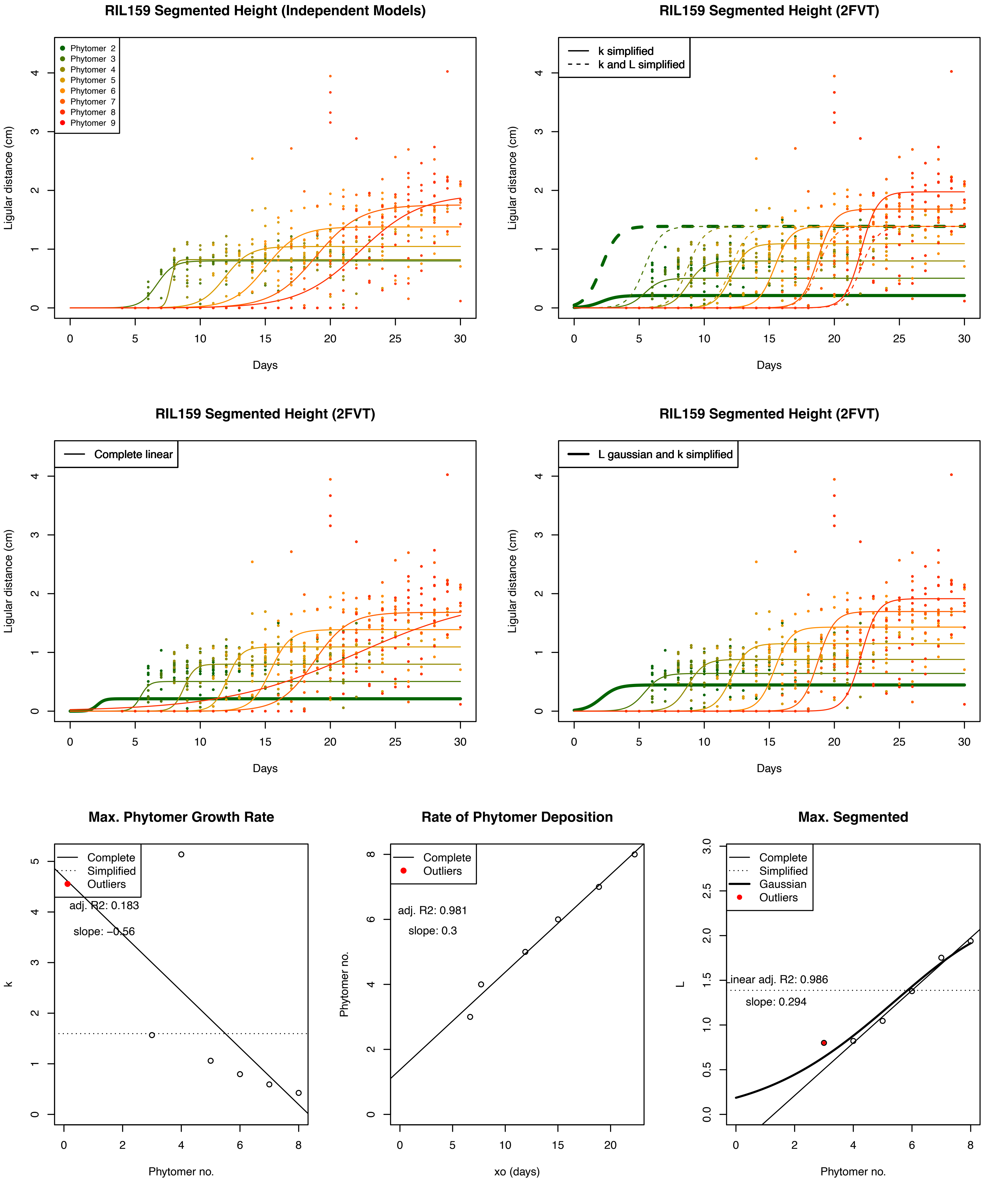
RIL159 Segmented Height (Independent Models)
Phytomer 2
Phytomer 3
Phytomer 4
Phytomer 5
Phytomer 6
Phytomer 7
Phytomer 8
Phytomer 9
Ligular distance (cm)
Days
RIL159 Segmented Height (2FVT)
k simplified
k and L simplified
Ligular distance (cm)
Days
RIL159 Segmented Height (2FVT)
Complete linear
Ligular distance (cm)
Days
RIL159 Segmented Height (2FVT)
L gaussian and k simplified
Ligular distance (cm)
Days
Max. Phytomer Growth Rate
Complete
Simplified
Outliers
adj. R2: 0.183
slope: −0.56
k
Phytomer no.
Rate of Phytomer Deposition
Complete
Outliers
adj. R2: 0.981
slope: 0.3
Phytomer no.
xo (days)
Max. Segmented
Complete
Simplified
Gaussian
Outliers
Linear adj. R2: 0.986
slope: 0.294
L
Phytomer no.
