## Supplementary figures and images for "Wheels within wheels, Nested Function-value Traits as a Tool for Modeling Ontogeny"

### Supplementary Figure 1

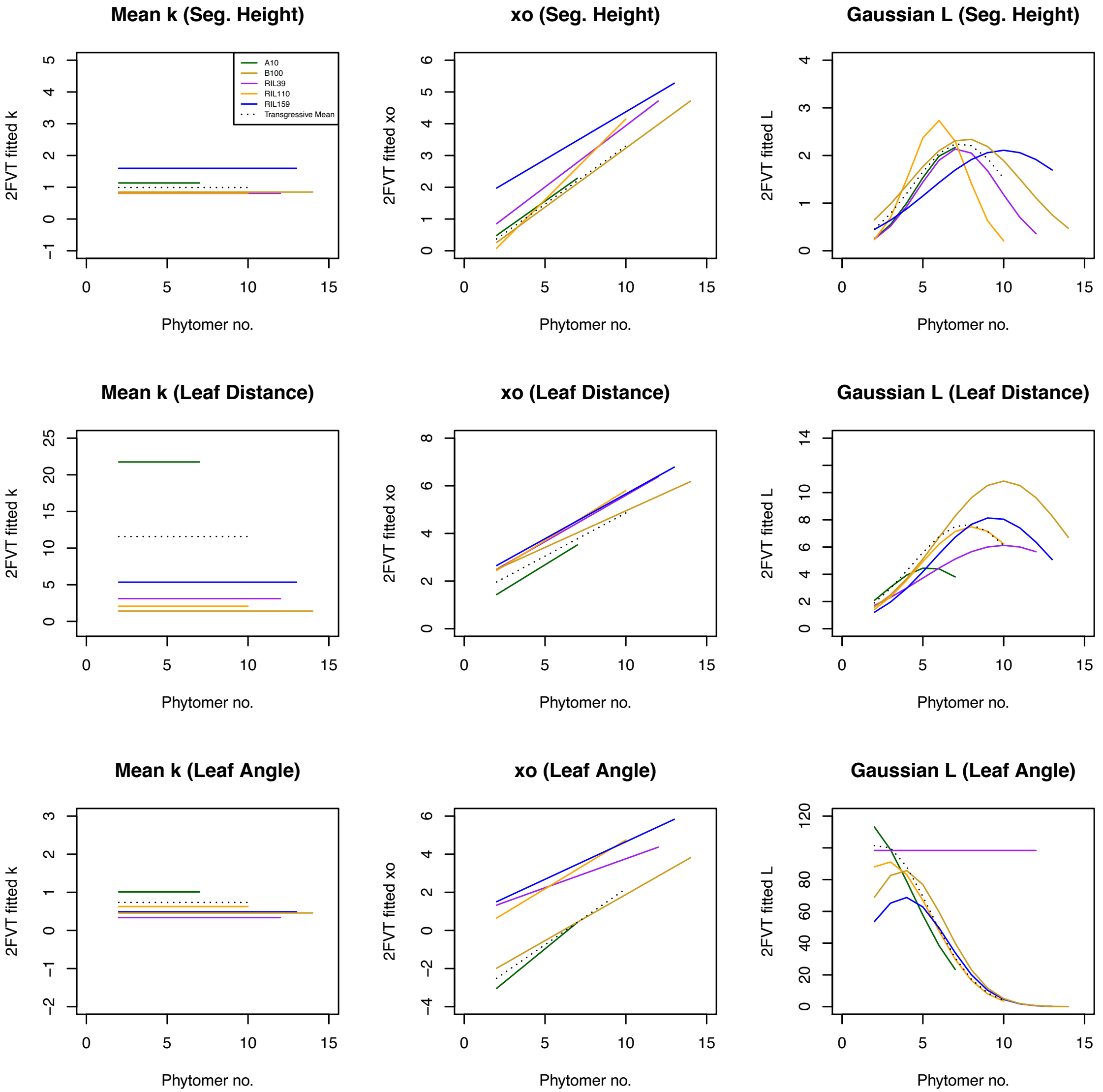
